# Targeting REV1 Exposes Context-Dependent Therapeutic Responses in Colorectal Cancer

**DOI:** 10.64898/2026.11.04.764315

**Authors:** Vihit Gupta, Madison Guyette, Sombodhi Bhattacharya, Joshua A. Victor, Naveen Kothandaraman, Katherine Boyle, Jon P. Girard, Jiyong Hong, Jon Ramsey, Pei Zhou, Nimrat Chatterjee

**Author notes:** To whom correspondence should be addressed: Nimrat Chatterjee, M.Sc., Ph.D. Equal contribution (Vihit Gupta, Madison Guyette, Sombodhi Bhattacharya).

## Abstract

Translesion synthesis (TLS) promotes DNA damage tolerance and contributes to chemotherapy resistance by enabling replication across damaged DNA templates. REV1, a central TLS regulator, is a potential therapeutic target to enhance the efficacy of genotoxic chemotherapy. Here, we investigated the therapeutic consequences of pharmacologic and genetic REV1 targeting in colorectal cancer (CRC) and evaluated the sensitivity of patient-derived CRC organoids to the REV1 inhibitor JH-RE-06. JH-RE-06 enhanced oxaliplatin sensitivity in HCT116 cells and 5-fluorouracil (5-FU) sensitivity in HT-29 cells, whereas reciprocal combinations produced limited sensitization, indicating treatment-and cellular context-dependent responses. siRNA-mediated REV1 depletion similarly enhanced chemotherapy sensitivity, supporting REV1 involvement in these responses. MNNG sensitivity assays and MLH1 expression analyses did not reveal consistent treatment-associated changes in mismatch repair function, while targeted HPRT1 sequencing identified no detectable sequence alterations in the region examined. Chemosensitization was not accompanied by consistent increases in caspase-3/7 activation under the conditions tested. REV1 inhibition also caused LC3-II accumulation and, after prolonged inhibitor exposure, reduced β-tubulin abundance, indicating additional autophagy-associated and cytoskeletal responses. In two patient-derived CRC organoid models, JH-RE-06 exhibited single-agent growth-inhibitory activity, with mean IC50 values of 2.01 and 1.4 µM, respectively. Together, these findings demonstrate context-dependent chemosensitization and organoid growth inhibition following REV1 targeting, suggesting that its therapeutic consequences may extend beyond canonical TLS-mediated DNA damage tolerance.

**SIGNIFICANCE STATEMENT:** Targeting REV1 enhances sensitivity to specific chemotherapeutic agents in colorectal cancer cells and inhibits growth of patient-derived colorectal cancer organoids. REV1 inhibition is also accompanied by LC3-II accumulation and reduced β-tubulin abundance, revealing cellular responses beyond conventional measures of DNA damage and mutagenesis. These findings identify treatment-dependent therapeutic vulnerabilities and suggest that the consequences of REV1 targeting may extend beyond its established role in translesion synthesis, supporting further investigation of REV1-directed therapeutic strategies in colorectal cancer.

**CLINICAL TRANSLATION RELEVANCE:** Chemotherapy resistance remains a major obstacle to effective treatment of colorectal cancer (CRC). This study identifies REV1 as a potential therapeutic target, showing that pharmacologic and genetic REV1 targeting enhances oxaliplatin sensitivity in HCT116 cells and 5-fluorouracil (5-FU) sensitivity in HT-29 cells, while reciprocal combinations show limited additional sensitization. Furthermore, patient-derived CRC organoids demonstrate single-agent sensitivity to REV1 inhibition at low-micromolar concentrations, supporting its activity across distinct tumor-derived models. REV1 inhibition is also associated with LC3-II accumulation and reduced β-tubulin abundance, suggesting additional cellular responses that may influence therapeutic outcomes. Together, these findings provide a preclinical rationale for investigating REV1-directed therapies in CRC and underscore the importance of identifying tumor-specific determinants of chemotherapy response and sensitivity to REV1 inhibition.

**Graphical abstract:** 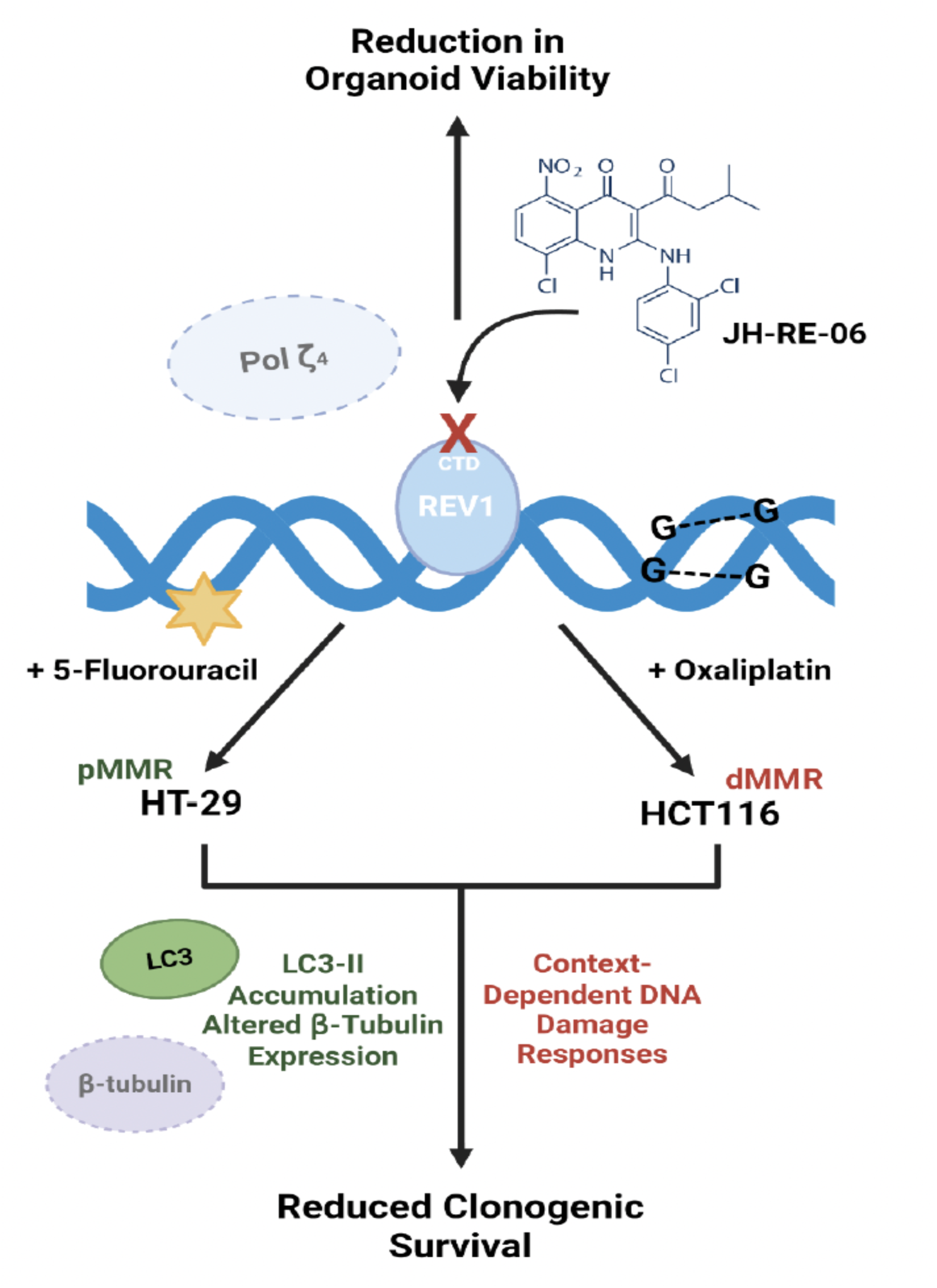

**Graphical abstract. Schematic representation of REV1 targeting and treatment-dependent chemosensitization in colorectal cancer** Pharmacologic inhibition of REV1 using JH-RE-06 disrupts its interaction with polymerase ζ (POL ζ) and selectively enhances chemotherapy sensitivity in colorectal cancer (CRC) cells. REV1 inhibition increases sensitivity to 5-fluorouracil (5-FU) in mismatch repair-proficient (pMMR) HT-29 cells and to oxaliplatin in mismatch repair-deficient (dMMR) HCT116 cells, resulting in reduced clonogenic survival. REV1 inhibition is accompanied by increased LC3-II abundance and reduced β-tubulin expression, indicating autophagy-associated and cytoskeletal changes. Assays of mismatch repair-associated responses, HPRT mutagenesis, DNA damage, and apoptosis did not identify a consistent mechanism explaining the differential chemosensitization. JH-RE-06 also exhibits single-agent growth-inhibitory activity in patient-derived CRC organoids. Together, these findings highlight treatment-dependent therapeutic vulnerabilities associated with REV1 targeting in CRC.

## INTRODUCTION

Translesion synthesis (TLS) is a specialized DNA damage tolerance pathway that enables DNA replication to proceed across lesions that would otherwise impede replication fork progression [1–3]. Unlike DNA repair pathways that remove or resolve DNA lesions, TLS employs specialized DNA polymerases to bypass damaged templates, allowing replication to continue despite persistent DNA damage [4, 5]. Although TLS promotes replication continuity under genotoxic stress, the relatively low fidelity of several TLS polymerases can also contribute to mutagenesis and genomic instability. In cancer cells, TLS-mediated lesion bypass can promote survival following genotoxic chemotherapy, while treatment-associated mutagenesis may contribute to the emergence of resistant tumor populations [1, 2, 6]. REV1/polymerase ζ-dependent TLS has therefore emerged as a potential therapeutic target to disrupt DNA damage tolerance and enhance chemotherapy efficacy [7–11].

REV1 is a central TLS regulator, with both deoxycytidyl transferase activity and a scaffolding function that coordinates specialized DNA polymerases through protein– protein interactions [2, 4]. In particular, the REV1 C-terminal domain (CTD) mediates interactions with REV7, a regulatory subunit of DNA polymerase ζ, facilitating polymerase recruitment during lesion bypass. The small-molecule inhibitor JH-RE-06 binds the REV1 CTD and promotes REV1 dimerization, thereby disrupting the REV1–REV7 interaction and impairing mutagenic TLS [7]. JH-RE-06 enhances cisplatin-induced cytotoxicity and suppresses treatment-associated mutagenesis in preclinical models, establishing the therapeutic potential of targeting REV1-dependent polymerase interactions [7, 10]. Subsequent investigations showed that combining JH-RE-06 with cisplatin also promotes senescence-associated responses, suggesting that REV1 inhibition may trigger cellular processes beyond conventional apoptotic responses to DNA damage [8].

Colorectal cancer (CRC) represents an important setting for investigating REV1-directed therapeutic strategies because intrinsic and acquired chemotherapy resistance remain major barriers to effective treatment [12, 13]. Fluoropyrimidine-based regimens, frequently combined with oxaliplatin or irinotecan, remain central to systemic CRC therapy [14–16]. However, these agents exert their cytotoxic effects through distinct molecular mechanisms. Oxaliplatin generates platinum-associated DNA lesions, predominantly intrastrand crosslinks, that interfere with DNA replication and transcription [17, 18]. In contrast, 5-fluorouracil (5-FU) disrupts thymidylate synthesis and nucleotide metabolism and can induce DNA-and RNA-associated stress [19, 20]. Resistance to these agents involves multiple mechanisms, including altered drug metabolism, DNA damage processing, and adaptive stress responses. Given the distinct molecular consequences of oxaliplatin and 5-FU treatment, determining whether REV1 inhibition enhances sensitivity to both agents, or instead produces treatment-dependent responses, is important for evaluating its therapeutic potential in CRC. Previous investigations have shown that REV1 targeting can alter cellular responses to genotoxic therapy beyond its established role in mutagenic TLS [8, 21]. Pharmacologic REV1 inhibition with JH-RE-06 enhances senescence-associated responses following cisplatin treatment [8], whereas subsequent studies identified autophagy-associated responses following REV1 inhibition and demonstrated that this approach does not necessarily enhance radiosensitivity in HCT116 colorectal cancer cells [21]. Furthermore, previous investigations demonstrated that genetic or pharmacologic REV1 inhibition represses β-tubulin expression at both the transcriptional and translational levels, accompanied by G2/M cell-cycle arrest [22]. Together, these findings suggest that the biological and therapeutic consequences of REV1 inhibition may depend on the nature of the genotoxic challenge. However, whether REV1 targeting produces distinct chemosensitization responses across CRC cellular backgrounds and chemotherapy agents remains incompletely understood. Furthermore, the relationship between REV1 inhibition, DNA damage-associated responses, autophagy-related changes, and microtubule-associated proteins warrants investigation. Evaluating REV1 inhibitor activity in patient-derived CRC organoids may also help establish their therapeutic potential across heterogeneous tumor-derived models.

In this study, we investigated the therapeutic and cellular consequences of pharmacologic and genetic REV1 targeting in CRC models. We demonstrate that REV1 inhibition preferentially enhances oxaliplatin sensitivity in HCT116 cells and 5-FU sensitivity in HT-29 cells, whereas reciprocal combinations produce limited additional sensitization. Complementary genetic experiments support REV1 involvement in these treatment-dependent responses. We further examined mismatch repair function, targeted HPRT1 sequence alterations, DNA damage-associated responses, and caspase-3/7 activation to characterize cellular outcomes, specifically DNA damage, damage response, and apoptosis-associated mechanisms accompanying REV1 inhibition. Notably, REV1 targeting was associated with LC3-II accumulation and reduced β-tubulin abundance under the respective treatment conditions, suggesting engagement of additional molecular mechanisms beyond canonical TLS-mediated DNA damage tolerance. Finally, JH-RE-06 showed single-agent growth-inhibitory activity in two clinically distinct patient-derived CRC organoid models, including one derived from a treatment-naïve tumor and another from a previously treated tumor that had progressed following FOLFOX (folinic acid (leucovorin), 5-fluorouracil, and oxaliplatin)-based therapy. Together, these findings identify treatment-and cellular context-dependent responses to REV1 targeting and support further investigation of REV1-directed therapeutic strategies in CRC.

## METHODS

### Cell Culture

Human colorectal cancer cell lines HCT116 (RRID:CVCL_0291) and HT-29 (RRID:CVCL_0320) were obtained from the American Type Culture Collection (ATCC, Manassas, VA, USA) and maintained in McCoy’s 5A medium (Gibco, Thermo Fisher Scientific; Cat. No. 16600082) supplemented with 10% fetal bovine serum (FBS; Thermo Fisher Scientific) and 1% penicillin–streptomycin (Thermo Fisher Scientific). Mouse areolar fibroblastic cell line L-WRN (stably expressing human Wnt-3a, R-spondin 3, and Noggin) was obtained from the American Type Culture Collection (ATCC, Manassas, VA, USA) and maintained in DMEM (high glucose, Corning Cat. No. 10-013-CV) supplemented with 10% heat-inactivated fetal bovine serum (Omega Scientific, Cat. No. 110617) and 1% Antibiotic/Antimycotic solution (Gibco, Thermo Scientific, Cat. No. 15420-062). This cell line was used to prepare conditioned media containing Wnt-3a, R-spondin 3, and Noggin as recommended by the ATCC.

Wild-type (WT) and *REV1* knockout (KO) mouse embryonic fibroblasts (MEFs) were kindly provided by Dr. Neil de Wind’s laboratory. The *REV1*KO MEFs harbor a deletion of exon 10 of the mouse *Rev1* gene, resulting in a frameshift that disrupts the REV1 C-terminal domain (CTD), which mediates interactions with translesion synthesis-associated proteins, including REV7 [23]. Thus, this genetic model complements pharmacologic disruption of REV1 CTD-mediated protein interactions by JH-RE-06. MEFs were maintained in Dulbecco’s modified Eagle medium (DMEM; Gibco, Thermo Fisher Scientific) containing 4.5 g/L D-glucose and supplemented with 10% FBS and 1% penicillin–streptomycin.

All cell lines were maintained at 37 °C in a humidified atmosphere containing 5% CO₂ and were routinely passaged using 0.25% trypsin–EDTA (Gibco, Thermo Fisher Scientific Cat. No. 25200072).

### Clonogenic Survival Assay (CSA)

Cells were seeded at a density of 1,000 cells per well in six-well plates containing 3 mL of complete culture medium. After 24 hours, cells were treated with JH-RE-06 (1 µM), oxaliplatin (1 µM; Selleckchem, Cat. No. S1209), or 5-fluorouracil (5-FU; 1 µM; Selleckchem, Cat. No. S1224), either individually or in the indicated combinations. Untreated cells served as controls. Drug exposure was maintained for either 24 or 72 hours, depending on the experimental design. For the 72-hour treatment conditions, the original drug-containing medium remained unchanged throughout the exposure period, without drug replenishment. Following treatment, drug-containing medium was replaced with fresh complete medium, and cells were maintained at 37 °C in a humidified atmosphere containing 5% CO₂ to allow colony formation. Once visible colonies had developed in untreated control wells, the medium was aspirated, and colonies were fixed with methanol and stained with 1 mL of Coomassie Brilliant Blue solution per well. Colonies containing more than 50 cells were counted microscopically. Relative clonogenic survival was calculated by normalizing colony counts in treated wells to their corresponding untreated controls.

### siRNA-Mediated Knockdown of REV1

HCT116 and HT-29 cells were seeded at a density of 50,000 cells per well in 12-well plates. Twenty-four hours after seeding, cells were transfected with a pool of three REV1-targeting small interfering RNAs (siRNAs; Sigma-Aldrich), each at a final concentration of 10 nM (30 nM total), using Lipofectamine RNAiMAX (Thermo Fisher Scientific, Cat. No. 13778030) and Opti-MEM (Gibco, Thermo Fisher Scientific, Cat. No. 31985062). Transfections were performed in serum-free McCoy’s 5A medium for 4 hours, after which the transfection medium was replaced with fresh complete McCoy’s 5A medium. Mock-transfected cells received the transfection reagents without siRNA.

Following a 24-hour recovery period, cells underwent a second round of transfection using the same siRNA pool and conditions. After an additional 24-hour recovery period, cells were enzymatically dissociated and replated in six-well plates for clonogenic survival assays, as described above. Total RNA was isolated using the RNeasy Mini Kit (Qiagen, Cat. No. 74104), and REV1 transcript abundance was assessed by reverse transcription–quantitative PCR (RT-qPCR) to determine knockdown efficiency. The sequences of the three REV1-targeting siRNAs are provided in the accompanying siRNA sequence in **Table 1**.

**Table 1.**
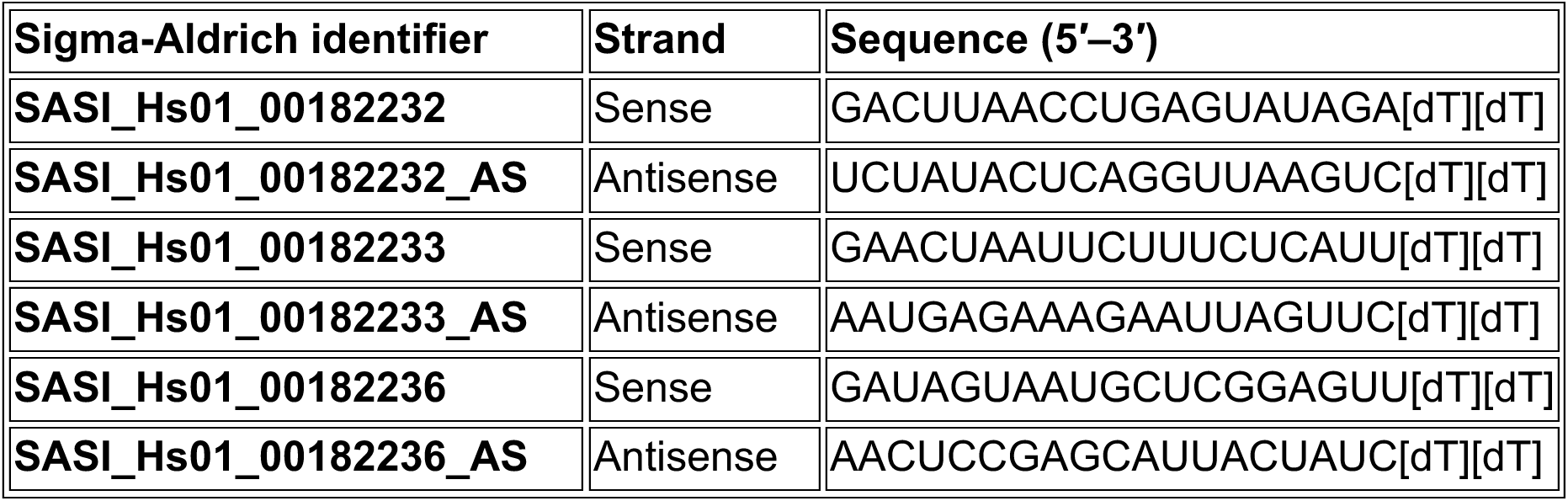
siRNA sequences used for siREV1-mediated knockdown.

### Western Blot Analysis

Cells were lysed in radioimmunoprecipitation assay (RIPA) buffer (G-Biosciences, Cat. No. 786-490) supplemented with protease and phosphatase inhibitor cocktails (Roche, Cat. No. 4906845001). Total protein concentrations were determined using the Micro BCA Protein Assay Kit (Pierce, Thermo Fisher Scientific, Cat. No. 23235). Protein lysates were diluted with a mix of 2× Laemmli sample buffer and β-mercaptoethanol (β-ME) and heated at 95–100 °C for 5 minutes before separation by sodium dodecyl sulfate– polyacrylamide gel electrophoresis (SDS-PAGE). Proteins were transferred to polyvinylidene difluoride (PVDF) membranes using the Mini Blot Module (Thermo Fisher Scientific). Membranes were blocked with 5% nonfat dry milk (Carnation, Nestlé). Primary and secondary antibodies were diluted in 5% bovine serum albumin (BSA) prepared in Tris-buffered saline containing 0.01% Tween-20 (TBST).

Membranes were incubated overnight at 4 °C with the following primary antibodies: β-actin (Thermo Fisher Scientific, Cat. No. MA1-744; RRID: AB_2223496; 1:1,000), MLH1 (Cell Signaling Technology, clone 4C9C7; RRID: AB_2145615; 1:250), LC3-I/II (Thermo Fisher Scientific, Cat. No. PA1-16931; RRID: AB_2137583; 1:250), SQSTM1/p62 (Thermo Fisher Scientific, Cat. No. PA5-20839; RRID: AB_11157045; 1:250), β-tubulin (Cell Signaling Technology, Cat. No. 2146; RRID:AB_2210545; 1:500), and GAPDH (Cell Signaling Technology, clone 14C10; RRID:AB_561053; 1:500).

Following four washes with TBST, membranes were incubated with the appropriate fluorescent secondary antibodies, IRDye 800CW goat anti-rabbit IgG (LI-COR Biosciences, Cat. No. 926-32211; RRID: AB_2651127) or IRDye 680RD goat anti-mouse IgG (LI-COR Biosciences, Cat. No. 926-68070; RRID: AB_10956588), diluted 1:20,000. Immunoreactive bands were visualized using a LI-COR Odyssey imaging system. Band intensities were quantified using Image Studio software (LI-COR Biosciences; RRID:SCR_015795). Relative protein abundance was determined by normalizing the intensity of each target protein band to its corresponding β-actin or GAPDH loading control, followed by normalization to the appropriate experimental control. GAPDH was used for normalization of LC3-II and β-tubulin abundance in the indicated experiments. For LC3 analyses, the LC3-II band was quantified.

### Reverse Transcription–Quantitative PCR (RT-qPCR)

Total RNA was isolated using the RNeasy Mini Kit (Qiagen) according to the manufacturer’s instructions. RNA concentrations were determined using a NanoDrop spectrophotometer (Thermo Fisher Scientific). REV1 transcript abundance was quantified by one-step reverse transcription–quantitative PCR (RT-qPCR) using the iTaq Universal SYBR Green One-Step Kit (Bio-Rad, Cat. No. 1725150), with 10 ng of total RNA per reaction.

Reverse transcription was performed at 50 °C for 10 minutes, followed by an initial denaturation step at 95 °C for 1 minute and 40 amplification cycles consisting of 95 °C for 15 seconds and 60 °C for 1 minute.

Relative REV1 transcript abundance was determined using the comparative threshold cycle (2^−ΔΔCt) method, with GAPDH as the endogenous reference gene and mock-transfected cells as the calibrator. **Table 2** provides the primer sequences used for REV1 and GAPDH amplification.

**Table 2.**
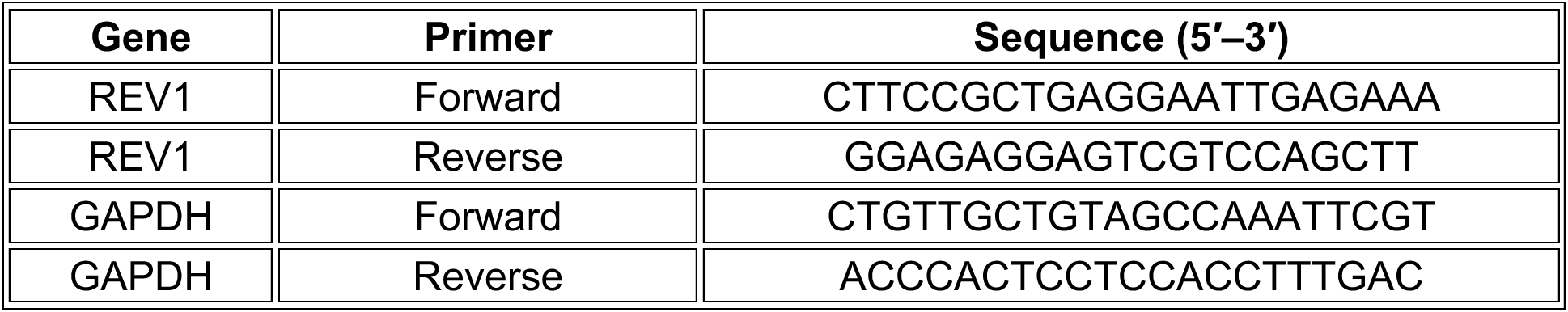
Primer sequences used for RT-qPCR.

### Targeted HPRT1 Sequencing Analysis

Following the indicated treatments, HCT116 and HT-29 cells were harvested, and genomic DNA was isolated using Qiagen reagents. Exon 6 of the hypoxanthine phosphoribosyltransferase 1 (*HPRT1*) gene was amplified by polymerase chain reaction (PCR) using 20 ng of genomic DNA, Platinum SuperFi II PCR Master Mix (Thermo Fisher Scientific, Cat. No. 12368010), and gene-specific primers synthesized by Integrated DNA Technologies (IDT), as listed in **Table 3**. PCR amplification was performed using a MiniAmp Plus Thermal Cycler (Applied Biosystems, Thermo Fisher Scientific) under the following cycling conditions: initial denaturation at 98 °C for 30 seconds; 35 cycles of denaturation at 98 °C for 10 seconds, annealing at 60 °C for 10 seconds, and extension at 72 °C for 1 minute; followed by a final extension at 72 °C for 5 minutes and a hold at 4 °C. PCR products were purified using the QIAquick PCR Purification Kit (Qiagen) according to the manufacturer’s instructions and submitted to Azenta Life Sciences (GENEWIZ) for Sanger sequencing.

**Table 3.**
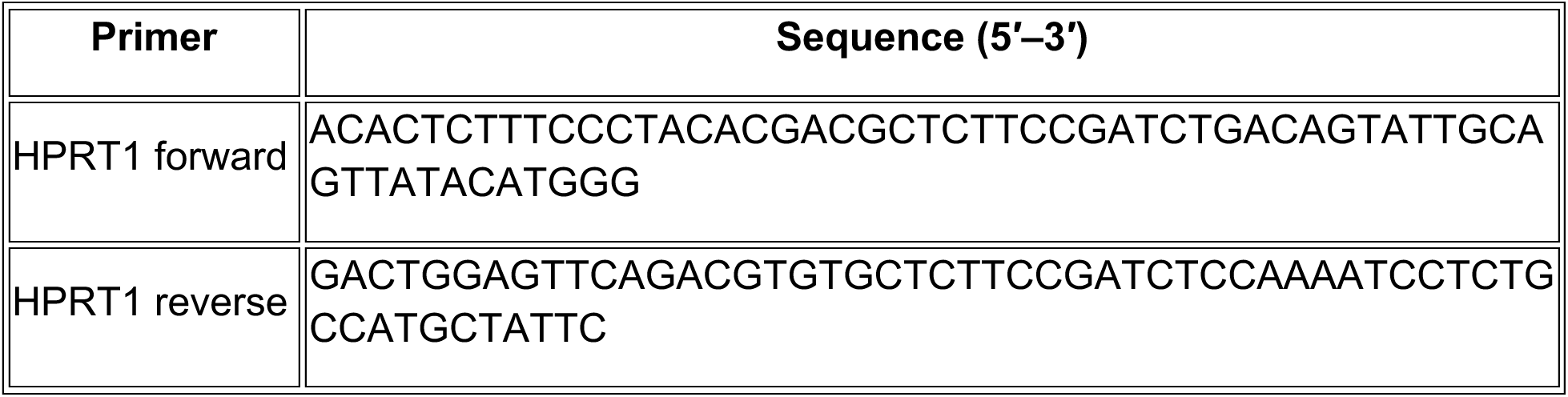
Primers used for HPRT1 exon 6 amplification.

Sequencing data obtained from PCR products representing pooled cell populations were analyzed using Geneious software (version 2022.0.2; RRID: SCR_010519). Sequences were compared with the *HPRT1* reference sequence obtained from the Ensembl genome database (gene ID: ENSG00000165704; genome assembly: GRCh37/hg19; RRID: SCR_002344), focusing on exon 6 of transcript 1. Detectable nucleotide substitutions or other sequence alterations within the amplified region were assessed by comparing sequencing results across experimental conditions. Because sequencing was performed on PCR products derived from pooled cell populations, this analysis was restricted to detectable sequence alterations within the targeted *HPRT1* region.

### Alkaline Comet Assay

HCT116 and HT-29 cells were seeded in six-well plates and treated for 24 hours with JH-RE-06 (1 µM), either alone or in combination with oxaliplatin (1 µM) in HCT116 cells or 5-fluorouracil (5-FU; 1 µM) in HT-29 cells. Following treatment, cells were washed with phosphate-buffered saline (PBS), detached using 0.25% trypsin–EDTA (Thermo Fisher Scientific, Cat. No. 25200072), collected by centrifugation, and washed with PBS. Alkaline single-cell gel electrophoresis was performed using selected reagents from the Comet SCGE Assay Kit (Enzo Life Sciences, Cat. No. ADI-900-166). Glass microscope slides (Fisherbrand, Cat. No. 12-550-15) were precoated with 2% agarose and air-dried overnight. The supplied low-melting-point agarose was melted at 65 °C and allowed to cool while remaining molten. A 50 µL aliquot of the cell preparation was mixed with 500 µL of low-melting-point agarose, and 200 µL of the resulting suspension was immediately applied to each precoated slide. Samples were covered with coverslips and placed on a prechilled ice pack for approximately 9 minutes to allow the agarose to solidify.

Following solidification, coverslips were carefully removed, and slides were incubated in prechilled lysis buffer supplied with the kit for 2 hours at 4 °C. Slides were subsequently transferred to an electrophoresis chamber containing chilled alkaline electrophoresis buffer (300 mM NaOH and 1 mM EDTA, pH approximately 13) and incubated for 15 minutes to allow DNA unwinding. Electrophoresis was performed at a set current of 280 mA for 30 minutes. Following electrophoresis, slides were washed twice with 0.4 M Tris buffer and once with ultrapure water. DNA was stained overnight with 1× CYGREEN dye (Enzo Life Sciences).

Comet images were acquired using a Nikon Eclipse Ts2R microscope equipped with a FITC fluorescence filter (excitation/emission approximately 489/515 nm). Images were analyzed using the OpenComet plugin for ImageJ (RRID:SCR_003070) [24]. DNA migration was quantified as the percentage of DNA in the comet tail (% tail DNA), with 100 comets evaluated per treatment condition. Statistical analyses were performed using GraphPad Prism 10 (RRID:SCR_002798).

### Fluorescence Microscopy and Micronucleus Quantification

HCT116 and HT-29 colorectal cancer cells were seeded at a density of 10,000 cells per well in eight-well chamber slides (Corning, Cat. No. 354118) and allowed to adhere for 24 hours. Cells were subsequently treated with JH-RE-06 (1 µM), oxaliplatin (1 µM), or 5-fluorouracil (5-FU; 1 µM), individually or in the indicated combinations, for 24 hours. Following treatment, culture medium was removed, and cells were mounted using ProLong™ Diamond Antifade Mountant with DAPI (Invitrogen Cat. No. P36962) to visualize nuclear DNA. Fluorescence images were acquired using a Marianas microscopy system (Intelligent Imaging Innovations, Denver, CO, USA) comprising a Zeiss Axio Observer 7 microscope equipped with a motorized XY stage, an Okolab stage-top incubation chamber, an X-Cite NOVEM-L LED illumination system (Excelitas), and an ORCA-Fusion BT sCMOS camera (Hamamatsu). Images were acquired using a 63×/1.4 NA Plan-Apochromat oil-immersion objective with differential interference contrast (DIC) capability. Micronuclei were identified as discrete DAPI-positive extranuclear structures distinct from the primary nucleus and manually quantified. 100 cells per treatment condition were evaluated across three independent biological experiments.

### Caspase Glo 3/7 Assay

Caspase-3/7 activity was assessed using the Caspase-Glo 3/7 Assay (Promega, Cat. No. G8981) according to the manufacturer’s instructions. HCT116 and HT-29 cells were seeded at a density of 5,000 cells per well in white-walled, clear-bottom 96-well plates and allowed to adhere for 24 hours under standard culture conditions (37 °C, 5% CO₂).

Cells were subsequently treated with JH-RE-06 (1 µM), oxaliplatin (1 µM), or 5-fluorouracil (5-FU; 1 µM), individually or in the indicated combinations, for 24 hours. Treatments were performed in triplicate, with untreated cells serving as negative controls.

Following treatment, the Caspase-Glo 3/7 reagent (Promega) was added to each well. Plates were incubated at room temperature for 2 hours to allow generation of the luminescent signal associated with caspase-3/7 activity. Luminescence was measured using a BioTek Neo2 multimode plate reader. Raw luminescence values were normalized to the mean signal of the corresponding untreated controls, and relative caspase-3/7 activity was analyzed using GraphPad Prism 10 (RRID:SCR_002798).

### Patient-Derived Colorectal Cancer Organoid Culture and Maintenance

Human colorectal adenocarcinoma organoid models were obtained from the National Cancer Institute Patient-Derived Models Repository (NCI-PDMR). The models included 116655-072-R-V1 (116655), derived from a primary mucinous adenocarcinoma from a 63-year-old male patient (NCI-PDMR, lot CK11220), and 919269-233-R5-V5 (919269), derived from rectosigmoid primary tumor colorectal adenocarcinoma from a 66-year-old male patient (NCI-PDMR, lot CK9997). Other clinical and molecular characteristics of these two patient-derived colorectal cancer organoids are in **Table 4**.

**Table 4.** Clinical and molecular characteristics of patient-derived colorectal cancer organoid models.

| **Characteristic** | **Organoid 116655** | **Organoid 919269** |
|---|---|---|
| NCI PDMR model identifier | 116655-072-R-V1-organoid | 919269-233-R5-V5-organoid |
| Source | NCI PDMR | NCI PDMR |
| Model derivation | Patient-derived xenograft-derived organoid | Patient-derived xenograft-derived organoid |
| Biological sex | Male | Male |
| Age at diagnosis | 63 years | 65 years |
| Date of diagnosis | February 2019 | April 2018 |
| Histological subtype | Mucinous colon adenocarcinoma | Rectosigmoid adenocarcinoma |
| Microsatellite instability status | MSI-high (reported) | Not confirmed |
| Mismatch repair status | Not independently confirmed | PDMR annotation pMMR-; interpretation pending |
| Reported molecular alterations | APC | APC, TCF7L2, TP53 |
| Prior systemic therapy | Treatment-naïve | Bevacizumab + FOLFOX |
| Number of treatment cycles | 0 | 12 |

Organoids were maintained and propagated in Media 6C/Colon 1B, prepared according to NCI-PDMR standard operating procedure SOP30101 (https://dctd.cancer.gov/drug-discovery-development/reagents-materials/pdmr/using-pdmr/sops). Organoids were cultured as hydrogel domes formed by inversion in Cultrex PathClear Reduced Growth Factor Basement Membrane Extract (BME), Type 2 (R&D Systems, Cat. No. 3533-005-02), using CellStar suspension culture plates (Greiner Bio-One, Cat. No. M9062). Cultures were maintained at 37 °C in a humidified incubator containing 5% CO₂.

For routine passaging, organoid-containing BME domes were incubated with Dispase II (1 mg/mL; Sigma-Aldrich) prepared in D-BSA solution consisting of high-glucose Dulbecco’s modified Eagle medium (DMEM; Gibco), penicillin–streptomycin (100 U/mL; Gibco), and 0.1% bovine serum albumin (BSA; Sigma-Aldrich/Millipore). Domes were incubated for 2 hours at 37 °C, with hydrogel disruption facilitated by gentle pipetting. Intact organoids were subsequently washed with D-BSA and collected by centrifugation at 300 × g for 4 minutes.

To generate smaller organoid fragments for passaging, organoid pellets were resuspended in undiluted TrypLE Express without phenol red (Gibco, Cat. No. 12-604-013) and incubated for 2 minutes at room temperature. Organoids were mechanically dissociated by repeated pipetting, with the extent of fragmentation monitored using an inverted microscope until fewer than 5% of organoids remained intact. TrypLE activity was quenched by adding 10 volumes of D-BSA, followed by centrifugation and removal of the supernatant. The resulting organoid fragments were resuspended in undiluted Cultrex BME and replated as hydrogel domes at a 1:5 passage ratio.

### Organoid Drug Sensitivity Assays

Patient-derived colorectal cancer organoids maintained in three-dimensional basement membrane extract (BME) domes were harvested by Dispase II digestion, as described above, and washed and resuspended in D-BSA solution. Organoids were size-selected by sequential filtration through 100 µm and 20 µm pore-size cell strainers (PluriSelect). Organoids measuring approximately 20–100 µm, retained on the 20 µm strainer, were recovered by inverting the strainer and washing with Dulbecco’s phosphate-buffered saline (DPBS). Size-selected organoids were counted and resuspended at a density of 10,000 organoids/mL in Media 6C/Colon 1B supplemented with 10% Cultrex BME. Aliquots of 50 µL, corresponding to approximately 500 organoids per well, were dispensed into black-walled 96-well culture plates (Costar, Corning).

JH-RE-06 was prepared from a 30 mM stock solution in dimethyl sulfoxide (DMSO), and a twofold serial dilution series was generated to obtain final treatment concentrations ranging from 0 to 100 µM. A 50 µL aliquot of each drug dilution was added to the corresponding organoid-containing wells, yielding a final culture volume of 100 µL per well. Treatments were performed in triplicate, with DMSO concentrations maintained consistently across experimental conditions. Organoids were exposed to JH-RE-06 for 48 hours at 37 °C in a humidified atmosphere containing 5% CO₂.

Following treatment, organoid cultures were imaged using a Cytation C10 Multimode Imager (Agilent Technologies) in brightfield and high-contrast imaging modes at 4× magnification. Organoid viability was subsequently assessed using the CellTiter-Glo 3D Cell Viability Assay (Promega, Cat. No. G9681) according to the manufacturer’s instructions. ATP-dependent luminescence was measured using a Synergy Neo2 Hybrid Multimode Reader (Agilent Technologies). Concentration–response data were analyzed using GraphPad Prism 10 (GraphPad Software; RRID:SCR_002798) and fitted by nonlinear least-squares regression to a four-parameter, variable-slope inhibition model to determine half-maximal inhibitory concentrations (IC50). IC50 values were determined independently for each of three biological replicate experiments per organoid model and summarized as the mean ± standard deviation.

For complementary morphological and viability assessment, drug-exposed organoids were stained using the Viability/Cytotoxicity Assay Kit (Proteintech Cat. No. PF00007) according to the manufacturer’s instructions. Organoids were imaged using the Cytation C10 in brightfield and widefield fluorescence modes at 4× magnification to assess morphology and visualize calcein-AM fluorescence in viable cells and ethidium homodimer-2 (EthD-2) fluorescence in membrane-compromised cells. Z-stack images were acquired at 5 µm intervals across a total depth of 50 µm, yielding 11 optical sections per field. Images were processed using Gen5 software (Agilent Technologies) to generate maximum-intensity projections.

### Statistical Analysis

Statistical analyses were performed using GraphPad Prism 10 (GraphPad Software; RRID:SCR_002798). Data are presented as mean ± standard error of the mean (SEM) or standard deviation (SD), as indicated in the corresponding figure legends. For comparisons involving multiple treatment groups, ordinary one-way analysis of variance (ANOVA) was performed, followed by Tukey’s or Holm–Šídák’s multiple-comparisons test, as appropriate. Comparisons between two independent groups were performed using unpaired Student’s t-tests. The statistical tests applied to individual experiments are specified in the corresponding figure legends.

For patient-derived colorectal cancer organoid drug-sensitivity studies, concentration– response data were fitted by nonlinear regression using a four-parameter, variable-slope inhibition model. IC50 values were determined independently for each of three biological replicate experiments per organoid model and summarized as mean ± SD.

Statistical significance was defined as ***P*** < 0.05. Exact P values, where available, and the number of independent biological replicates are reported in the corresponding figure legends. Technical replicates were distinguished from independent biological experiments in the statistical analyses.

## RESULTS

### Pharmacologic REV1 inhibition produces treatment-dependent chemosensitization in colorectal cancer cells

To determine whether pharmacologic REV1 inhibition alters chemotherapy sensitivity in colorectal cancer (CRC), we examined two CRC cell lines with distinct mismatch repair (MMR) backgrounds: MMR-deficient HCT116 cells and MMR-proficient HT-29 cells. Cells were treated with the REV1 inhibitor JH-RE-06 in combination with oxaliplatin or 5-fluorouracil (5-FU), and long-term survival was assessed using clonogenic survival assays following 24-or 72-hour drug exposure.

In HCT116 cells, JH-RE-06 significantly enhanced oxaliplatin-associated suppression of clonogenic survival following both 24-and 72-hour treatment **(Fig. 1A).** This response was treatment-specific, as JH-RE-06 did not significantly enhance the effects of 5-FU at either treatment duration **(Supplementary Fig. S1A-B)**. In HT-29 cells, JH-RE-06 significantly enhanced 5-FU-associated suppression of clonogenic survival after 24-hour treatment, whereas no significant additional effect was detected after 72-hour exposure **(Fig. 1B).** Conversely, JH-RE-06 did not significantly enhance oxaliplatin-associated suppression of clonogenic survival in HT-29 cells at either treatment duration of 24 or 72 hours (**Supplementary Fig. S1D-E**), suggesting increased exposure of cells to oxaliplatin still did not enhance cytotoxicity in the HT-29 cells.

**Figure 1.**
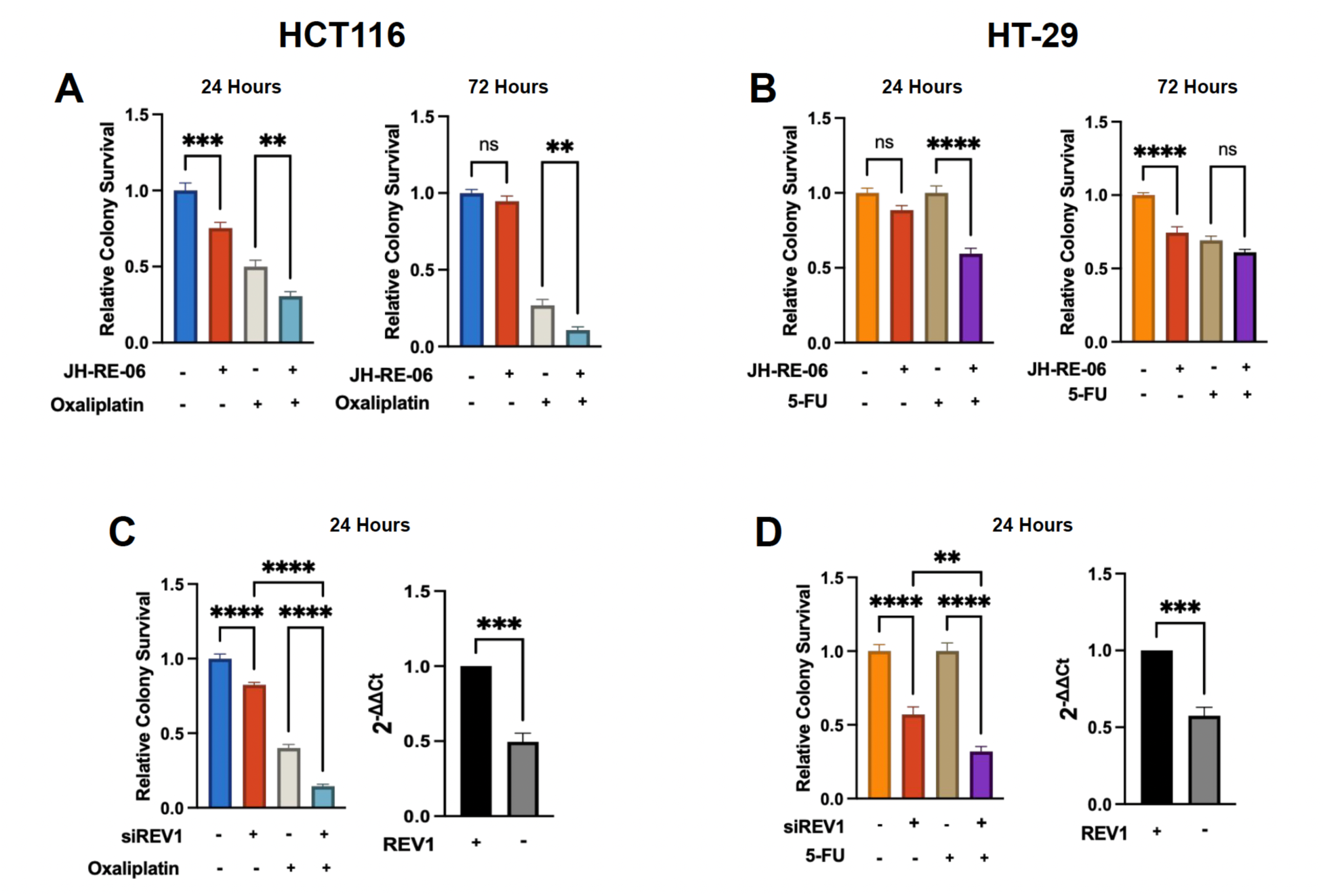
Pharmacologic and genetic REV1 targeting enhances chemotherapy sensitivity in a treatment-dependent manner in colorectal cancer cells. (A, **B)** Relative clonogenic survival of HCT116 (A) and HT-29 (B) cells following treatment with JH-RE-06 (1 µM), oxaliplatin (1 µM; HCT116), or 5-fluorouracil (5-FU; 1 µM; HT-29), individually or in combination, for 24 or 72 hours. **(C, D)** Relative clonogenic survival following siRNA-mediated REV1 depletion in HCT116 cells treated with oxaliplatin (C) and HT-29 cells treated with 5-FU (D) for 24 hours. REV1 transcript abundance was assessed by RT-qPCR and expressed relative to mock-transfected controls using the 2^−ΔΔCt method. Clonogenic survival was normalized to the corresponding untreated or mock-transfected control and is presented as mean ± SEM. Clonogenic survival comparisons were analyzed using ordinary one-way ANOVA followed by Tukey’s multiple-comparisons test; REV1 transcript expression was analyzed using an unpaired t-test. Statistical significance is indicated in the panels (ns, not significant; \*\****P*** < 0.01; \*\*\****P*** < 0.001; \*\*\*\****P*** < 0.0001). N=3 biological replicates, with 9 total technical replicates.

Because oxaliplatin and 5-FU did not enhance JH-RE-06-dependent cytotoxicity in the HT-29 and HCT116 cells, respectively, we asked whether the addition of the responsive treatments, 5-FU in HT-29 and oxaliplatin in the HCT116 cells, would re-establish the suppression of clonogenic survival in these models. We therefore examined whether JH-RE-06 enhanced the effects of combined 5-FU and oxaliplatin treatment in either cell line and observed no statistically significant reduction of clonogenic survival in either cell line even with 72-hour exposure **(Supplementary Fig. S1C and D).** Together, these findings demonstrate treatment-specific chemosensitization by pharmacologic REV1 inhibition, with enhanced oxaliplatin sensitivity in HCT116 cells and 5-FU sensitivity in HT-29 cells following 24-hour exposure. The absence of additional sensitization with the triple combination of JH-RE-06, oxaliplatin, and 5-FU suggests that REV1 inhibition does not simply amplify the cytotoxicity of every DNA-damaging treatment; rather, REV1’s contribution to chemotherapy tolerance may depend on the specific DNA lesion, replication-associated stress, and the cellular pathway engaged by each treatment. These observations suggest that the therapeutic consequences of REV1 inhibition depend on cellular background, chemotherapeutic agent, and treatment duration.

### siREV1-mediated depletion enhances chemotherapy sensitivity in colorectal cancer cells

To investigate whether genetic targeting of REV1 produces effects consistent with pharmacologic inhibition, we depleted REV1 using a pool of three REV1-targeting small interfering RNAs (siRNAs). RT-qPCR analysis confirmed an approximately 50% reduction in REV1 transcript abundance in both HCT116 and HT-29 cells (**Fig. 1C,D**). In HCT116 cells, REV1 depletion reduced clonogenic survival relative to mock-transfected controls and significantly enhanced oxaliplatin-associated suppression of colony formation (**Fig. 1C**). Similarly, in HT-29 cells, REV1 depletion reduced clonogenic survival and significantly enhanced the reduction in clonogenic survival following 5-FU treatment (**Fig. 1D**). These findings provide complementary genetic evidence supporting REV1 involvement in the chemotherapy responses observed following JH-RE-06 treatment.

### REV1-dependent chemosensitization is not associated with detectable changes in mismatch repair-associated responses

The treatment-dependent chemosensitization observed following REV1 targeting prompted us to investigate whether mismatch repair (MMR) function contributes to the differential chemotherapy sensitivity of HCT116 and HT-29 cells, which have documented MMR-deficient and MMR-proficient backgrounds, respectively [25, 26]. We examined cellular sensitivity to N-methyl-N′-nitro-N-nitrosoguanidine (MNNG), an alkylating agent commonly used to assess functional MMR. MMR-proficient cells recognize and process MNNG-induced DNA lesions, triggering cytotoxic responses, whereas MMR-deficient cells frequently exhibit resistance to MNNG [25, 27–29].

We first examined our established REV1-knockout mouse embryonic fibroblast (MEF) model [7, 23]. MNNG significantly reduced clonogenic survival in wild-type MEFs, consistent with functional MMR, whereas REV1-knockout MEFs did not exhibit a significant reduction under the same conditions (**Fig. 2A**). This differential response suggests that REV1 loss may compromise MMR-associated processing of MNNG-induced lesions or promote alternative mechanisms of MNNG resistance, identifying a potentially important relationship between REV1 and the cellular response to alkylating DNA damage.

**Figure 2.**
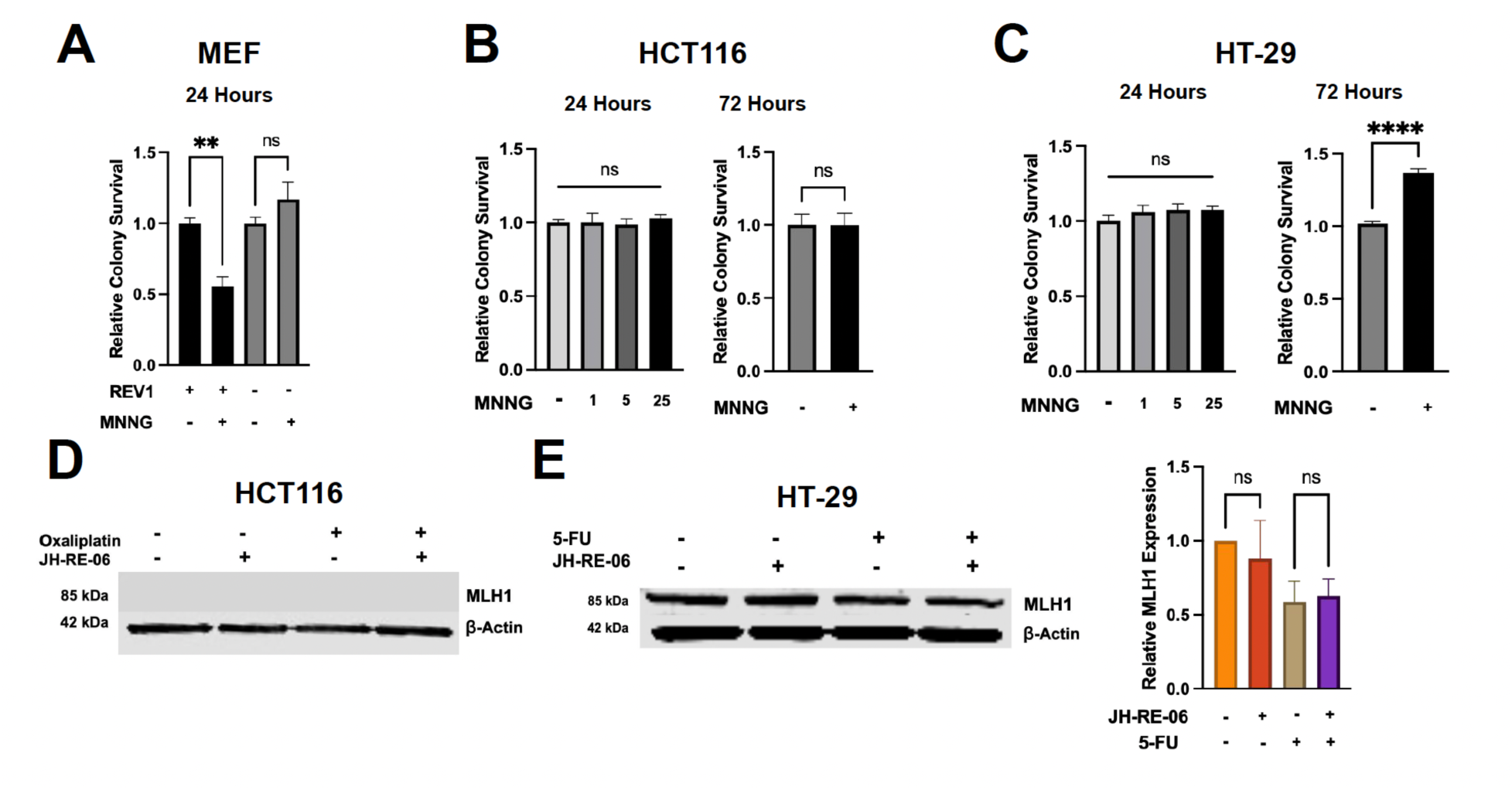
Assessment of mismatch repair-associated responses following REV1 targeting. **(A)** Relative clonogenic survival of wild-type and REV1-knockout mouse embryonic fibroblasts (MEFs) following exposure to 1 µM MNNG for 24 hours. **(B-C)** Relative clonogenic survival of HCT116 **(B)** and HT-29 **(C)** cells following exposure to the DNA-alkylating agent N-methyl-N′-nitro-N-nitrosoguanidine (MNNG). Cells were exposed to the indicated MNNG concentrations (1, 5, or 25 µM) for 24 hours, with additional survival measurements following the indicated 1 µM treatment for 72 hours. **(D–E)** Representative immunoblots examining MLH1 protein abundance in HCT116 cells treated with oxaliplatin with or without JH-RE-06 **(D)** and HT-29 cells treated with 5-FU with or without JH-RE-06 **(E)**. β-Actin was used as a loading control. Quantification of MLH1 abundance relative to β-actin is shown for HT-29 cells in **(E)**. Unless otherwise indicated, oxaliplatin, 5-FU, and JH-RE-06 were used at 1 µM. Quantitative data are presented as mean ± SEM. Statistical comparisons were performed using ordinary one-way ANOVA with the indicated multiple-comparisons tests or unpaired t-tests for two-group comparisons. Statistical significance is indicated in the panels (ns, not significant; \*\****P*** < 0.01; \*\*\*\****P*** < 0.0001). N=3 biological replicates, with 9 total technical replicates.

We next evaluated MNNG sensitivity in CRC cells. In HCT116 cells, MNNG did not significantly alter clonogenic survival following either 24-or 72-hour exposure (**Fig. 2B**), consistent with their established MMR-deficient phenotype. Surprisingly, HT-29 cells, despite their documented MMR-proficient background, also exhibited resistance to MNNG. No significant reduction in clonogenic survival was observed following 24-hour exposure to the tested MNNG concentrations, while 72-hour exposure to 1 µM MNNG resulted in a significant increase in relative clonogenic survival (**Fig. 2C**). These findings suggest that sensitivity to the MNNG-induced alkylation damage does not uniformly reflect the reported MMR status of the CRC models under the experimental conditions examined.

To investigate whether REV1 inhibition and chemotherapy altered the expression of an essential MMR component, we assessed MLH1 protein abundance following treatment with JH-RE-06, chemotherapy, or their combination. MLH1 was not detectably expressed in HCT116 cells (**Fig. 2D**), consistent with their established MLH1-deficient background. In HT-29 cells, MLH1 protein abundance was numerically reduced following combined JH-RE-06 and 5-FU treatment; however, this difference was not statistically significant (**Fig. 2E**).

Together, these findings reveal an altered MNNG response following genetic REV1 loss in MEFs, the expected MNNG resistance in MMR-deficient HCT116 cells, and an unexpected MNNG-resistant phenotype in HT-29 cells despite their documented MMR proficiency. Because REV1 inhibition did not produce a statistically significant alteration in MLH1 abundance under the chemotherapy conditions examined, these results suggest that REV1-dependent chemosensitization in CRC cells may occur independently of detectable changes in the MMR-associated responses assessed here.

### REV1-dependent chemosensitization is not accompanied by increased HPRT mutagenesis

REV1 is best characterized for its role in TLS-mediated lesion bypass and mutagenesis [2]. We therefore asked whether the observed chemotherapy responses were associated with increased mutagenesis at the HPRT locus. This question was particularly relevant because REV1 targeting can affect both lesion tolerance and mutagenic DNA synthesis, processes that are related but not necessarily reflected by identical experimental endpoints. To examine treatment-associated mutagenesis at a defined genomic locus, we performed targeted Sanger sequencing of HPRT1 exon 6 using genomic DNA isolated from pooled HCT116 and HT-29 cell populations following the indicated treatments. Exon 6 of HPRT1 was selected for sequence analysis because mutations within this reg ion have been documented in HPRT mutational-spectrum studies involving 6-thioguanine-resistant cells [30]. Comparison of the amplified sequences with the corresponding HPRT1 reference sequence revealed no detectable treatment-associated nucleotide alterations within the analyzed region (**Supplementary Fig. 2**). These surprising results indicate that REV1 targeting did not produce detectable sequence changes at the examined HPRT1 locus under the experimental conditions and within the assay’s detection limits. Because this assay interrogates a defined genomic region, these results do not exclude treatment-associated mutational changes elsewhere in the genome.

### REV1-dependent chemosensitization is associated with distinct DNA damage responses without detectable caspase-3/7 activation

Translesion synthesis (TLS) can facilitate replication across damaged DNA templates, providing an important mechanism by which cells tolerate genotoxic stress [2]. Because REV1 coordinates TLS polymerase recruitment, its inhibition could compromise lesion bypass and alter the accumulation or processing of chemotherapy-induced DNA damage [1, 4]. Such changes might contribute to reduced clonogenic survival, either by increasing DNA damage burden or activating downstream cell death pathways. Having observed treatment-specific chemosensitization following REV1 targeting [7], we therefore investigated whether the effective chemotherapy combinations produced corresponding changes in DNA damage and apoptotic responses.

First, to examine DNA damage accumulation, we performed alkaline comet assays [31] in HCT116 cells treated with oxaliplatin and HT-29 cells treated with 5-fluorouracil (5-FU), with or without JH-RE-06. Both chemotherapeutic agents produced detectable DNA damage-associated responses under the conditions examined, as previously reported [32, 33] and as shown here (**Figs. 3A–B**). Interestingly, JH-RE-06 addition produced distinct effects in the two cellular backgrounds. In HCT116 cells, combined REV1 inhibition and oxaliplatin treatment increased comet-detected DNA damage relative to oxaliplatin alone, suggesting a small but significant increase in DNA damage burden. In contrast, in HT-29 cells, the combination of JH-RE-06 and 5-FU was associated with surprisingly reduced comet-detected DNA damage relative to 5-FU alone, suggesting no increase in DNA damage burden with the combination treatment. Thus, although both combinations enhanced chemotherapy sensitivity, they did not produce a uniform increase in DNA damage, suggesting that the magnitude of comet-detected damage alone does not consistently predict the observed loss of clonogenic survival. Further, the damage context, where oxaliplatin-and 5-FU-induced DNA damage and their consequent engagement of the REV1 TLS polymerase might explain these differences, as discussed in the discussion section.

**Figure 3.**
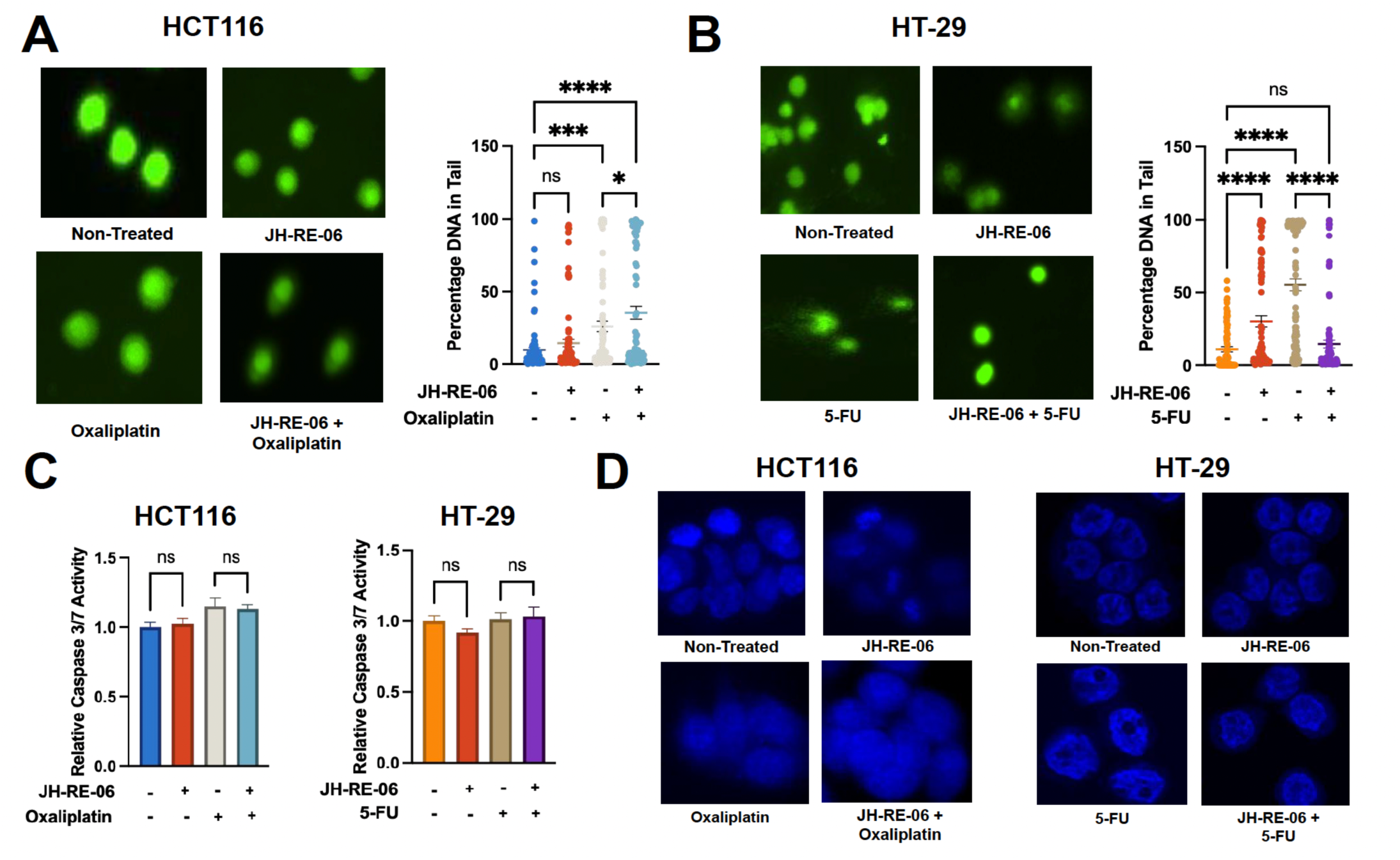
REV1-dependent chemosensitization is associated with distinct DNA damage responses without detectable caspase-3/7 activation. (A, **B)** Representative fluorescence micrographs and quantification of alkaline comet assays in HCT116 cells treated with oxaliplatin (A) and HT-29 cells treated with 5-fluorouracil (5-FU) (B), individually or in combination with JH-RE-06. DNA migration was quantified as the percentage of DNA in comet tails (% tail DNA). Individual points represent analyzed comets, and summary values are presented as mean ± SEM. **(C)** Relative caspase-3/7 activity in HCT116 and HT-29 cells following treatment with JH-RE-06, the corresponding chemotherapeutic agent, or their combination. Caspase activity was measured using the Caspase-Glo 3/7 assay and normalized to untreated controls. N=3 biological replicates with 9 technical replicates per biological replicate. **(D)** Representative DAPI-stained fluorescence micrographs of HCT116 and HT-29 cells following the indicated treatments, illustrating nuclear morphology and the assessment of micronucleus formation. N=3 biological replicates Unless otherwise indicated, JH-RE-06, oxaliplatin, and 5-FU were used at 1 µM for 24 hours. Comet assay comparisons were analyzed using one-way ANOVA followed by Tukey’s multiple-comparisons test. Statistical significance is indicated in the panels (ns, not significant; \**P* < 0.05; \*\*\**P* < 0.001; \*\*\*\**P* < 0.0001).

Second, we reasoned that persistent or inadequately resolved DNA damage can initiate apoptotic signaling [34]; so, we next investigated whether the reduced clonogenic survival was accompanied by activation of executioner caspases. We assessed caspase-3/7 activity following the indicated treatments [8, 35]; however, we detected no corresponding increase in the caspase-3/7 signal in either CRC cell line under the conditions examined (**Fig. 3C**). These findings suggest that enhanced chemotherapy sensitivity was not accompanied by detectable activation of these apoptotic effectors at the assessed time point. We additionally examined nuclear morphology and micronucleus formation as indicators of treatment-associated nuclear abnormalities [36]. We observed no detectable increase in micronucleus formation under the conditions tested, where the JH-RE-06 and combination of JH-RE-06 with either the oxaliplatin in HCT116, or with 5-FU in HT-29 cells (**Fig. 3D**). Collectively, these results show that REV1 targeting enhances chemotherapy sensitivity despite divergent comet-detected DNA damage responses and without corresponding increases in caspase-3/7 activation or micronucleus formation. These observations suggest that additional cellular responses may accompany the loss of clonogenic survival following REV1 inhibition.

### REV1 inhibition is associated with LC3-II accumulation and reduced β-tubulin abundance in CRC cells

Our previous investigation demonstrated that REV1 inhibition did not enhance radiosensitivity in HCT116 cells following ionizing radiation [21]. In contrast, the present study demonstrated enhanced oxaliplatin sensitivity following REV1 inhibition in the same cellular model, as evidenced by reduced clonogenic survival (**Fig. 1A and C**). Given the autophagy-associated responses identified in our earlier investigation [21], we examined whether REV1 inhibitor treatment altered autophagy-associated markers in CRC cells. Immunoblot analysis revealed increased LC3-II abundance following JH-RE-06 treatment in both HCT116 and HT-29 cells, including under the respective oxaliplatin and 5-FU combination conditions (**Figs. 4A–B**). In contrast, p62/SQSTM1 abundance remained relatively unchanged under the conditions examined. Thus, LC3-II accumulation was observed under chemotherapy combination conditions that also produced enhanced clonogenic sensitivity, identifying an autophagy-associated response accompanying REV1 inhibitor-mediated chemosensitization in both CRC models.

**Figure 4.**
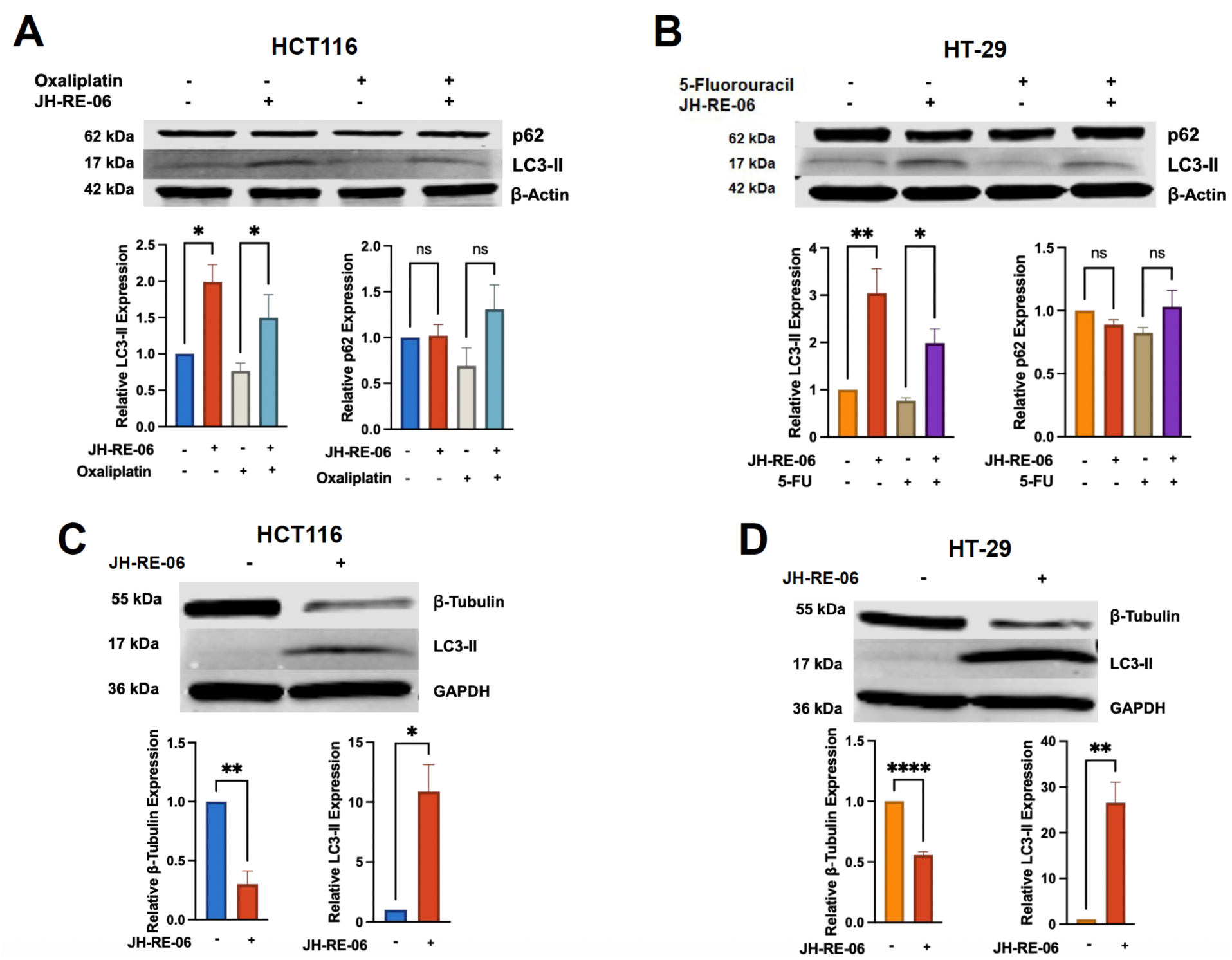
REV1 inhibition is associated with LC3-II accumulation and reduced β-tubulin abundance in colorectal cancer cells. (A–B) Representative immunoblots and corresponding densitometric analyses of LC3-II and p62/SQSTM1 abundance in HCT116 cells treated with oxaliplatin **(A)** and HT-29 cells treated with 5-fluorouracil (5-FU) **(B)**, with or without JH-RE-06. Cells were exposed to the indicated agents at 1 µM for 24 hours. β-Actin was used as the loading control, and relative protein abundance was quantified by densitometry. **(C–D)** Representative immunoblots and densitometric analyses of β-tubulin and LC3-II in HCT116 **(C)** and HT-29 **(D)** cells following single-agent treatment with 10 µM JH-RE-06 for 48 hours. GAPDH was included as a loading control. Quantitative immunoblot data are presented as mean ± SEM from three independent experiments. Statistical comparisons were performed using one-way ANOVA with Holm–Šídák multiple-comparisons testing (**A–B)** or an unpaired t-test **(C–D)**. Statistical significance is indicated in the panels (ns, not significant; \****P*** < 0.05; \*\*\****P*** < 0.001; \*\*\*\****P*** < 0.0001). N=3 biological replicates, with 9 total technical replicates.

Building on our previous findings linking REV1 inhibition to altered cell-cycle progression and reduced tubulin abundance [22], which demonstrates that REV1 loss disrupted tubulin expression transcriptionally and translationally, we asked whether REV1 targeting produced similar changes in colorectal cancer cells. Treatment of HCT116 and HT-29 cells with 10 µM JH-RE-06 for 48 hours resulted in significantly reduced β-tubulin abundance in both cell lines (**Figs. 4C–D**). Interestingly, analysis of the same protein samples also revealed pronounced LC3-II accumulation, consistent with the autophagy-associated changes observed under the shorter, lower-dose treatment conditions (**Figs. 4A–B**). Together, these observations demonstrate that prolonged REV1 inhibitor exposure is accompanied by reduced β-tubulin abundance and LC3-II accumulation, identifying additional cellular responses associated with pharmacological REV1 inhibition.

### Patient-derived colorectal cancer organoids representing distinct molecular and clinical backgrounds exhibit sensitivity to REV1 inhibition

The treatment-specific chemosensitization observed following REV1 targeting in HCT116 and HT-29 cells prompted us to investigate whether pharmacologic REV1 inhibition also exhibits activity in patient-derived colorectal cancer (CRC) models. Although established CRC cell lines provide valuable experimental systems for examining DNA damage tolerance and chemotherapy responses, they do not fully represent the molecular and clinical heterogeneity of colorectal malignancies [37]. In particular, differences in microsatellite instability, mismatch repair status, oncogenic alterations, and prior therapeutic exposure may influence cellular responses to DNA-damaging agents and targeted therapies [38]. Patient-derived organoid cultures, therefore, offer a complementary experimental platform for evaluating therapeutic responses in tumor models with clinically documented backgrounds. We therefore tested whether JH-RE-06 inhibits growth in patient-derived CRC organoids representing distinct molecular characteristics and treatment histories.

To address this question, we evaluated two independently derived CRC organoid models obtained from the National Cancer Institute Patient-Derived Models Repository (NCI PDMR). Both organoid cultures were established from patient-derived xenograft material and represented different colorectal tumor subtypes (**Table 4**). Organoid 116655-072-R-V1 originated from a treatment-naïve, 63-year-old male patient diagnosed with mucinous colon adenocarcinoma. We selected this model based on its reported microsatellite instability-high (MSI-H) classification and APC alterations, providing a tumor background of particular interest given the role of mismatch repair and DNA damage tolerance in chemotherapy responses. In contrast, organoid 919269-233-R5-V5 originated from a male patient diagnosed with rectosigmoid adenocarcinoma and harbored reported alterations involving APC, TCF7L2, and TP53. Importantly, this model’s clinical record documented disease progression after 12 cycles of bevacizumab plus FOLFOX, a chemotherapy regimen containing both 5-fluorouracil (5-FU) and oxaliplatin. Thus, the two models represented contrasting clinical treatment histories, including treatment-naïve disease and disease that had progressed following a standard chemotherapy-containing regimen.

The clinical background of organoid 919269 was particularly relevant to the present study because the preceding cell-line experiments identified treatment-dependent sensitization to oxaliplatin and 5-FU following REV1 targeting. However, the organoid experiments were designed to evaluate the intrinsic growth-inhibitory activity of JH-RE-06 as a single agent, rather than its ability to restore sensitivity to either chemotherapeutic agent. This distinction allowed us to assess whether pharmacologic REV1 inhibition is active in patient-derived CRC cultures independently of concurrent chemotherapy.

Both organoid models showed concentration-dependent reductions in viability-associated luminescence after 48 hours of exposure to JH-RE-06, as measured using CellTiter-Glo 3D (**Fig. 5A**). Analysis of three independent biological experiments demonstrated low-micromolar sensitivity in both models, with mean half-maximal inhibitory concentrations (IC50) of 2.10 ± 0.21 µM for organoid 116655 and 1.37 ± 0.38 µM for organoid 919269 (mean ± SD, n = 3 independent biological experiments per model; **Fig. 5C**). Although the previously treated model exhibited a numerically lower IC50, the limited number of organoid models does not permit conclusions regarding the relationship between molecular or clinical characteristics and drug sensitivity. These findings demonstrate that both patient-derived CRC organoid models were sensitive to pharmacologic REV1 inhibition at low-micromolar concentrations, despite differences in their reported molecular characteristics, histological subtypes, and clinical treatment histories.

**Figure 5.**
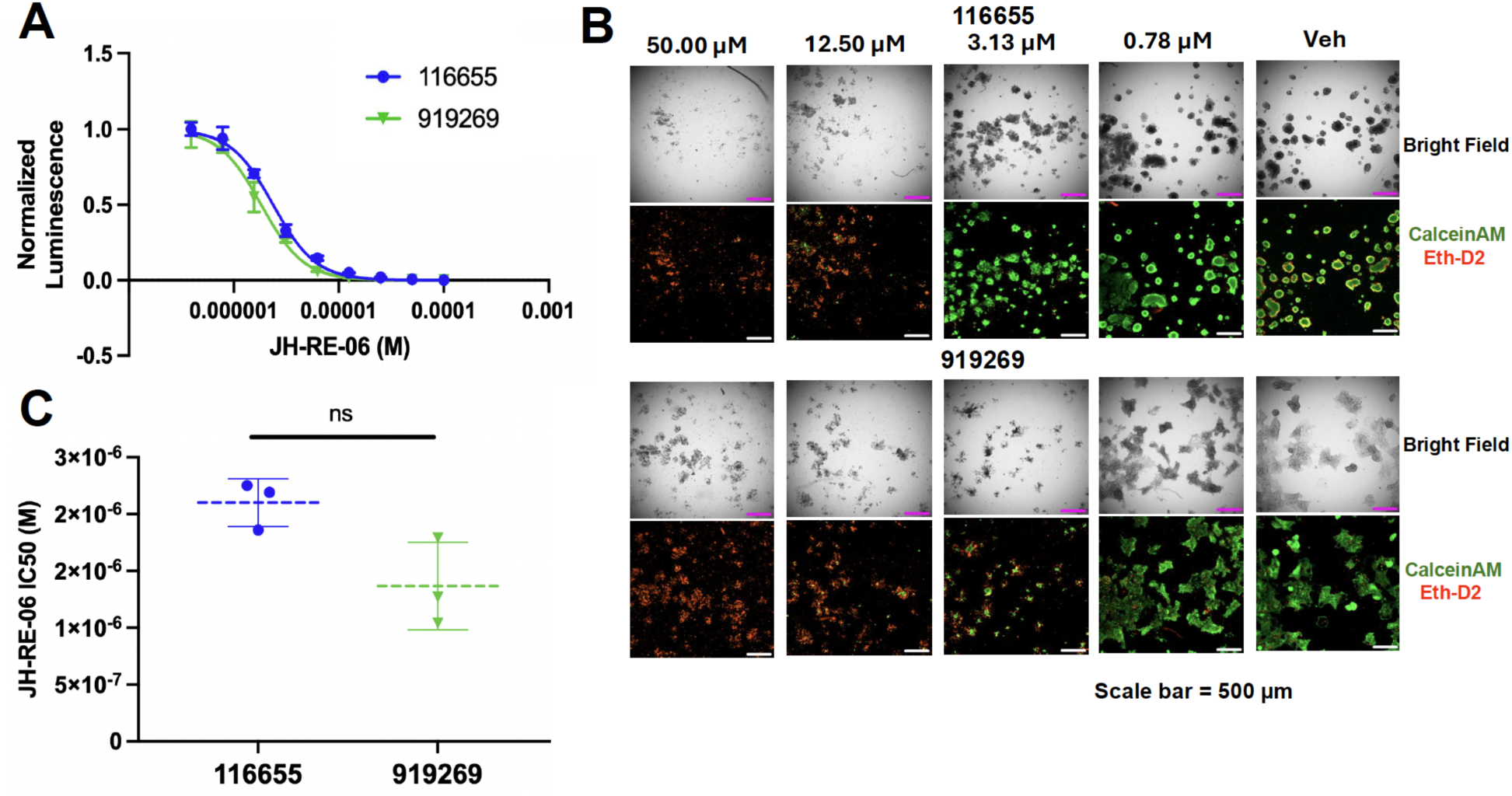
Patient-derived colorectal cancer organoids exhibit concentration-dependent sensitivity to pharmacologic REV1 inhibition. **(A)** Concentration–response curves showing the effects of JH-RE-06 on patient-derived colorectal cancer organoid models 116655-072-R-V1 and 919269-233-R5-V5 following 48 hours of treatment. Viability-associated luminescence was measured using CellTiter-Glo 3D and normalized to vehicle-treated controls. Concentration–response data were fitted using four-parameter variable-slope nonlinear regression, with the upper and lower plateaus constrained to 1 and 0, respectively. The pooled fitted IC50 values were 2.01 µM (95% CI, 1.898–2.124 µM) for organoid 116655 and 1.443 µM (95% CI, 1.363–1.527 µM) for organoid 919269. Data were obtained from three independent biological experiments. **(B)** Representative bright-field microscopy and Calcein-AM/EthD-2 live/dead fluorescence images of organoids following 48 hours of JH-RE-06 treatment. Calcein-AM fluorescence (green) identifies cells with intracellular esterase activity and intact membranes, whereas EthD-2 fluorescence (red) identifies cells with compromised plasma membrane integrity. Both organoid models exhibited concentration-dependent changes in morphology and viability-associated staining, including reduced Calcein-AM fluorescence and increased EthD-2 fluorescence at higher JH-RE-06 concentrations. Scale bars, 500 µm. **(C)** Half-maximal inhibitory concentration (IC50) values calculated independently from three biological experiments for each organoid model. Each symbol represents an independently determined IC50 value. Dashed horizontal lines indicate the mean, and error bars represent SD. Mean IC50 values were 2.10 ± 0.21 µM for organoid 116655 and 1.37 ± 0.38 µM for organoid 919269 (mean ± SD; n = 3 independent biological experiments per model).

To complement the quantitative viability measurements, we examined treatment-associated morphological changes and cell viability using bright-field microscopy and Calcein-AM/EthD-2 live/dead staining. Both organoid models exhibited concentration-dependent changes following JH-RE-06 treatment (**Fig. 5B**; **Supplementary Fig. S3**). At lower drug concentrations, organoids retained prominent Calcein-AM fluorescence, consistent with viable cells exhibiting intracellular esterase activity. In contrast, increasing JH-RE-06 concentrations were accompanied by diminished Calcein-AM staining and increased EthD-2-associated fluorescence, indicating a greater proportion of cells with compromised plasma membrane integrity. Bright-field microscopy also revealed changes in organoid morphology and the abundance of intact organoid structures at higher drug concentrations. These imaging observations complemented the concentration-dependent reductions in CellTiter-Glo 3D luminescence, providing independent qualitative evidence of reduced organoid viability following JH-RE-06 exposure.

Collectively, these results extend the observed growth-inhibitory activity of JH-RE-06 beyond established CRC cell lines and provide additional preclinical support for investigating REV1-directed therapeutic strategies in colorectal cancer. Notably, the activity observed in organoid 919269 demonstrates sensitivity to single-agent REV1 inhibition in a model derived from a patient with documented disease progression following FOLFOX-containing therapy. Although the limited number of organoid models precludes conclusions regarding molecular predictors of response, and the absence of chemotherapy combination experiments prevents assessment of resistance reversal, these findings establish a rationale for further evaluation of REV1 inhibition in larger, molecularly characterized patient-derived CRC organoid panels.

## DISCUSSION

Resistance to fluoropyrimidine-and platinum-based chemotherapy remains a major challenge in colorectal cancer (CRC) [39]. Translesion synthesis (TLS) enables replication across DNA lesions that would otherwise impede fork progression, but can also promote mutagenesis and treatment tolerance [1]. The REV1–DNA polymerase ζ (Polζ) axis is therefore an attractive target for disrupting DNA damage tolerance [2]. Here, we examined pharmacological REV1 inhibition with JH-RE-06 in CRC cell lines and patient-derived organoids. Our findings reveal chemotherapy-and cellular-context-dependent sensitization, distinct DNA damage-associated responses, and single-agent activity in clinically annotated organoid models.

REV1 functions as both a specialized DNA polymerase and a scaffold coordinating TLS polymerases [2]. Its interaction with REV7, a regulatory subunit of Polζ, supports lesion bypass, and previous studies demonstrated that disrupting REV1-dependent protein interactions can suppress mutagenic TLS and enhance responses to genotoxic agents [7, 8]. Our findings extend this therapeutic rationale to CRC while indicating that the consequences of REV1 targeting depend on the chemotherapeutic agent and cellular background. In HCT116 cells, JH-RE-06 enhanced oxaliplatin cytotoxicity, whereas HT-29 cells exhibited increased sensitivity to 5-fluorouracil (5-FU) under the tested conditions (**Fig. 1 and Supplementary Fig. 1**). Importantly, siRNA-mediated REV1 depletion similarly enhanced chemotherapy sensitivity, providing independent genetic evidence that REV1 contributes to chemotherapy tolerance in colorectal cancer cells. Notably, REV1-dependent chemosensitization was highly selective, with distinct responses to oxaliplatin and 5-FU in HCT116 and HT-29 cells, respectively, and no significant additional sensitization following the triple combination of JH-RE-06, oxaliplatin, and 5-FU under the conditions examined. These findings suggest that REV1’s contribution to chemotherapy tolerance is strongly influenced by the nature of treatment-induced DNA damage and the underlying cellular context, rather than reflecting a general requirement for REV1 in survival after genotoxic stress.

Elevated REV1 expression has been reported in colorectal tumors and associated with poorer patient outcomes, supporting its potential clinical relevance as a therapeutic target [40]. Nevertheless, our findings indicate that REV1-dependent chemotherapy tolerance is highly context-specific, raising important questions about how different forms of treatment-induced DNA damage engage REV1-mediated tolerance pathways. For example, following cellular uptake and aquation, oxaliplatin forms covalent platinum–DNA adducts, predominantly involving the N7 positions of purine bases, that interfere with DNA replication and transcription [17]. In contrast, 5-fluorouracil exerts its cytotoxic effects by inhibiting thymidylate synthase, perturbing nucleotide metabolism, and incorporating fluorinated metabolites into DNA and RNA [20, 41]. Consequently, their effects possibly engage overlapping but nonidentical damage-processing pathways.

We hypothesize that REV1-dependent TLS facilitates bypass of oxaliplatin-induced platinum–DNA lesions in HCT116 cells, and that JH-RE-06 disrupts REV1–Polζ interactions, compromising this damage-tolerance mechanism and reducing clonogenic survival after oxaliplatin treatment. In HT-29 cells, 5-FU-induced nucleotide pool imbalance and incorporation of uracil or fluorouracil into DNA may generate abasic sites and other replication-associated intermediates during DNA repair. REV1 could facilitate tolerance of these lesions through its catalytic activity and, particularly, its scaffolding function in recruiting Pol ζ to support translesion synthesis. Consistent with this possibility, REV1 and REV3 have been implicated in cellular tolerance to floxuridine [42], providing a mechanistic rationale for investigating REV1-dependent processing of 5-FU-associated DNA lesions in CRC cells. Although HCT116 and HT-29 cells differ in their established mismatch repair backgrounds, our MNNG response experiments and MLH1 expression analyses did not establish a direct relationship between REV1-dependent chemosensitization and mismatch repair status (**Fig. 2**). These observations suggest that additional DNA damage tolerance and repair pathways may contribute to the treatment-specific responses observed in these models.

In light of this, alkaline comet experiments further emphasized the context dependence of REV1 inhibition. Combined oxaliplatin and JH-RE-06 treatment increased comet tail DNA in HCT116 cells, consistent with increased alkali-detectable DNA damage or altered lesion processing (**Fig. 3A–B**). However, because the alkaline comet assay does not directly measure platinum–DNA adducts [43], the precise lesions underlying this response remain unresolved. Conversely, our reproducible data on combined 5-FU and JH-RE-06 treatment reduced comet tail DNA in HT-29 cells despite decreased clonogenic survival. This apparent discrepancy may reflect altered lesion processing rather than reduced cytotoxicity, as comet assays detect strand breaks and alkali-labile sites rather than the full spectrum of DNA lesions [43, 44]. Disruption of REV1-dependent lesion bypass may also alter replication-associated DNA processing, potentially reducing comet-detectable repair intermediates while compromising long-term cellular survival. Furthermore, enhanced chemosensitivity was not accompanied by increased caspase-3/7 activity or enhanced micronuclei formation under the conditions tested, suggesting that reduced clonogenic survival does not necessarily reflect increased executioner caspase activation. Delayed apoptosis or alternative consequences of replication-associated stress may therefore contribute to the observed treatment responses. Together, these findings suggest that REV1 inhibition produces distinct DNA damage-associated responses depending on the chemotherapeutic agent and cellular context.

REV1 inhibition was also associated with accumulation of the autophagy-related marker LC3-II (**Fig. 4**). During shorter treatments, p62 abundance did not exhibit a consistent corresponding change. Although these observations indicate altered autophagy-associated signaling or compartment abundance, LC3-II accumulation alone cannot establish increased autophagic flux, because it may reflect either enhanced autophagosome formation or impaired turnover [45]. Previous work from our laboratory connected REV1 targeting with altered cell-cycle progression and reduced tubulin expression [22]. Consistent with that report, prolonged JH-RE-06 exposure reduced β-tubulin abundance in both CRC cell lines and coincided with LC3-II accumulation. Together, these observations identify additional cellular responses to REV1 inhibition that warrant further investigation.

An important translational extension of this work was evaluating JH-RE-06 in patient-derived CRC organoids. Conventional cell lines cannot fully capture the molecular and clinical heterogeneity of colorectal malignancies, whereas patient-derived organoids offer complementary models that retain important tumor characteristics and enable evaluation of individualized therapeutic responses [38, 39]. Both organoid models exhibited concentration-dependent reductions in viability following single-agent JH-RE-06 treatment, with mean IC₅₀ values of 2.10 ± 0.21 µM for organoid 116655 and 1.37 ± 0.38 µM for organoid 919269 across three independent experiments (**Fig. 5**). Bright-field and Calcein-AM/EthD-2 imaging further demonstrated treatment-associated morphological changes and reduced viability at higher concentrations (**Fig. 5 and Supplementary Fig. 3**). Together, these findings establish reproducible low-micromolar activity of JH-RE-06 in patient-derived CRC organoids, extending its anticancer activity beyond conventional two-dimensional cell culture models.

Importantly, these organoids represented distinct clinical backgrounds (**Table 4**). Organoid 116655 originated from treatment-naïve mucinous colon adenocarcinoma, whereas organoid 919269 was derived from rectosigmoid adenocarcinoma in a patient whose disease progressed following bevacizumab-containing FOLFOX therapy. The latter model’s sensitivity to JH-RE-06 is particularly noteworthy given its prior exposure to a regimen containing 5-FU and oxaliplatin. Although single-agent activity does not establish reversal of FOLFOX resistance, it demonstrates that JH-RE-06 retains activity in a patient-derived model with a history of disease progression following standard chemotherapy. These findings support further investigation of REV1 inhibition in larger, molecularly and clinically annotated organoid panels, particularly in combination with established CRC chemotherapies, an approach supported by previous studies demonstrating the utility of patient-derived organoids in evaluating treatment responses [46].

Several limitations warrant consideration. Using two genetically distinct CRC cell lines precludes definitive identification of the molecular determinants underlying treatment-specific sensitization, and the DNA damage, caspase, and autophagy-associated measurements do not fully resolve the mechanisms involved. Furthermore, although the organoid experiments establish reproducible single-agent activity, additional studies are needed to determine *in vivo* efficacy and therapeutic benefit in combination with chemotherapy. Nevertheless, our findings provide compelling evidence that pharmacological disruption of REV1-dependent DNA damage tolerance can enhance chemotherapy sensitivity in a treatment-and cellular-context-dependent manner. Together with the low-micromolar activity observed in clinically distinct patient-derived CRC organoids, these results establish a strong preclinical rationale for further development of REV1-directed therapeutic strategies in colorectal cancer.

## Author Contributions

V.G. and M.G. ran colony assays, western blots, and caspase assays; S.B. ran COMET, immunofluorescence, qPCR assays, and helped analyze all the data; J.V. and N.K. ran HPRT mutagenesis assays; K.B. ran COMET assays and western blots; J.G. completed the micronuclei experiments; J.H. and P.Z. synthesized JH-RE-06; J.R. ran and supervised the organoid experiments; N.C. designed experiments, supervised research, analyzed data, and wrote the paper with help from all authors.

## Acknowledgments

We thank all Chatterjee Lab members for their valuable feedback; the UVM Cancer Center for access and assistance with shared resources; the UVM Cancer Center’s Organoid core for access to the organoid facility; and the Thali Lab for access to their microscope. This study is supported by the Nicole Ferland Award to Madison Guyette, the DUSRA award to Naveen Kothandaraman, the NIH R01CA279034 (to J.H. and P.Z.); a UVMCC pilot award and the NIGMS R35GM150992 to N.C.

---

**Supplementary Figure S1.**
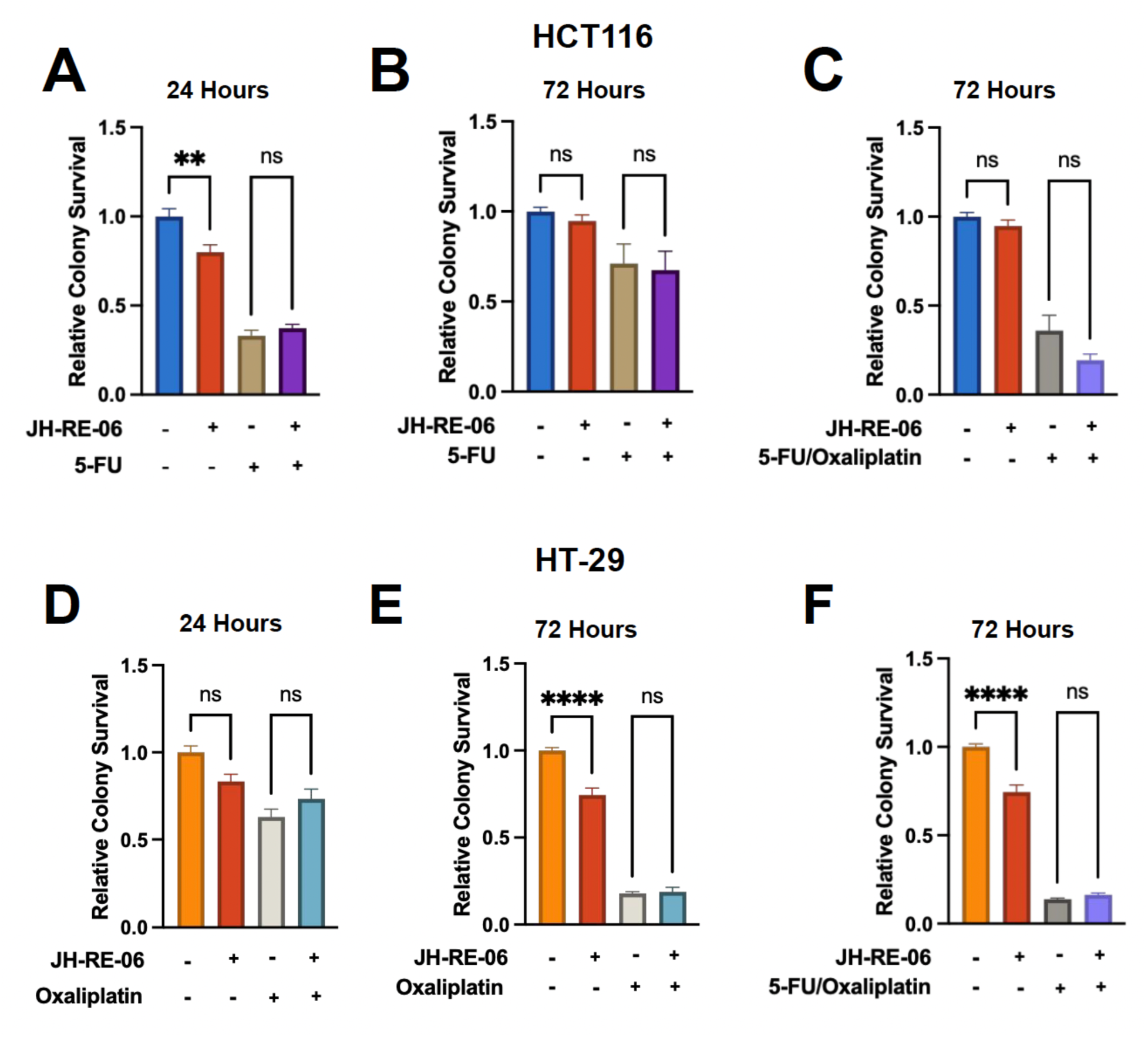
REV1-dependent CRC chemosensitization is specific across colorectal cancer cells. (A-C) Relative clonogenic survival of HCT116 cells following treatment with JH-RE-06 (1 µM) in combination with 5-fluorouracil (5-FU; 1 µM) for 24 or 72 hours, or with combined 5-FU and oxaliplatin (1 µM each) for 72 hours. **(D-F)** Relative clonogenic survival of HT-29 cells following treatment with JH-RE-06 (1 µM) in combination with oxaliplatin (1 µM) for 24 or 72 hours, or with combined 5-FU and oxaliplatin (1 µM each) for 72 hours. Cells were treated with individual agents or the indicated combinations, and clonogenic survival was assessed following drug removal and subsequent colony formation. Survival was normalized to the corresponding untreated control. Data are presented as mean ± SEM. Statistical comparisons were performed using ordinary one-way ANOVA followed by Tukey’s multiple-comparisons test. Significance annotations indicate the comparisons shown (ns, not significant; \*\****P*** < 0.01; \*\*\*\****P*** < 0.0001). Biological and technical replicate numbers will be specified following verification of the experimental records. N=3 biological replicates, with 9 total technical replicates.

**Supplementary Figure S2.**
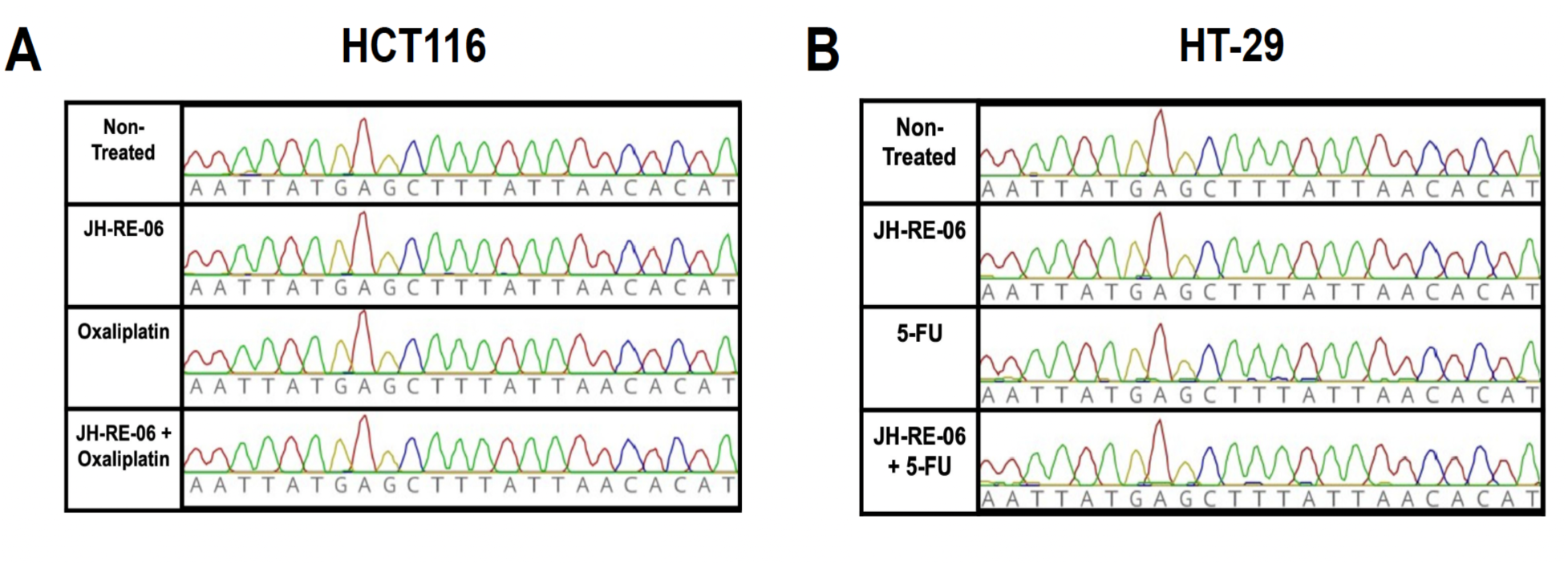
Targeted HPRT1 sequencing following REV1 inhibition and chemotherapy treatment in colorectal cancer cells. (A, **B)** Representative Sanger sequencing chromatograms of PCR-amplified *HPRT1* exon 6 from HCT116 (A) and HT-29 (B) cells following 24-hour treatment with JH-RE-06 (1 µM), oxaliplatin (1 µM; HCT116) or 5-fluorouracil (5-FU; 1 µM; HT-29), individually or in combination. Untreated cells served as controls. Genomic DNA was isolated from pooled cell populations, and amplified sequences were compared with the corresponding *HPRT1* reference sequence. We observed no detectable treatment-associated nucleotide differences within the analyzed region. N=3 biological replicates, with 9 total technical replicates.

**Supplementary Figure S3.**
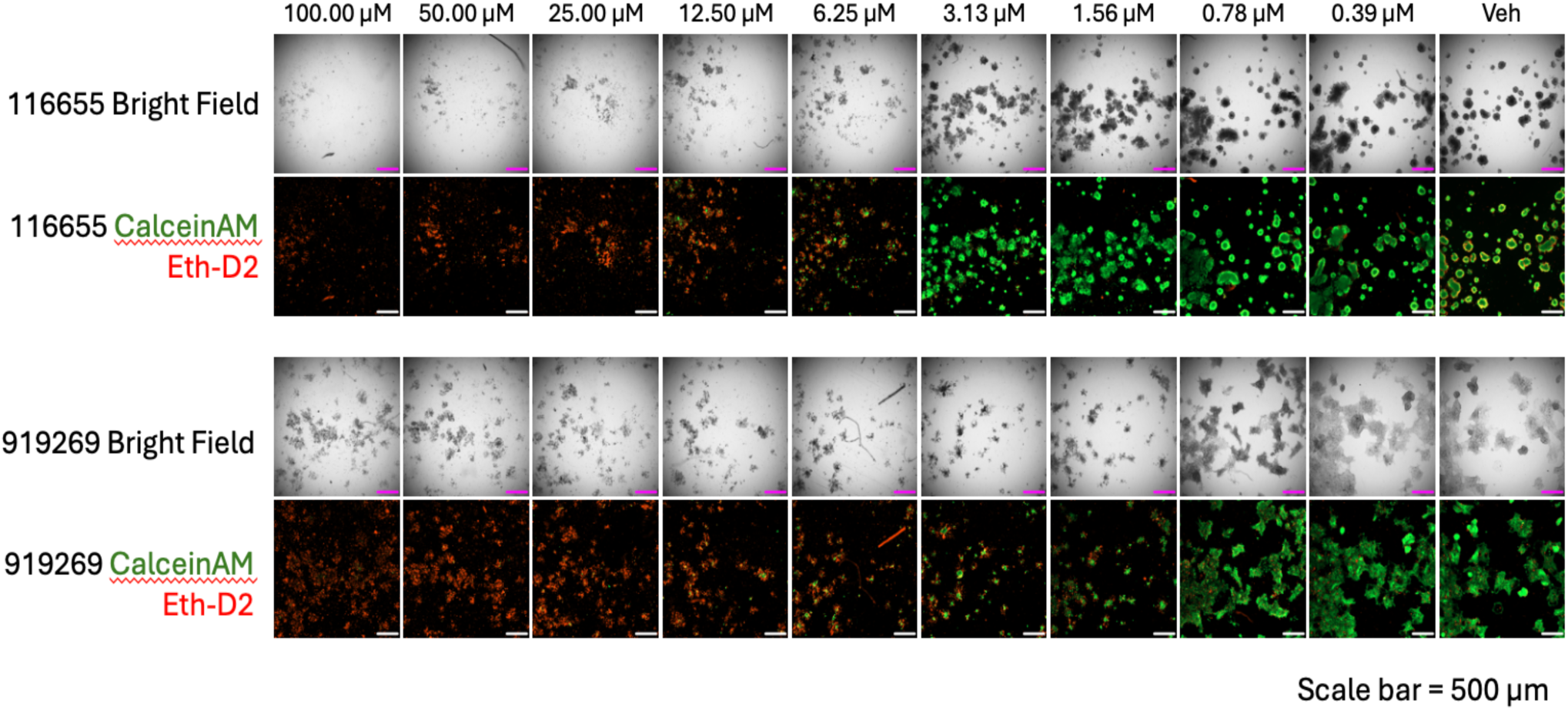
Concentration-dependent morphological and live/dead staining responses to JH-RE-06 in patient-derived colorectal cancer organoids. Representative bright-field and Calcein-AM/EthD-2 fluorescence images of patient-derived colorectal cancer organoid models 116655-072-R-V1 and 919269-233-R5-V5 following 48 hours of treatment with vehicle or the indicated JH-RE-06 concentrations (0.39–100 µM). Calcein-AM fluorescence (green) indicates viable cells with intracellular esterase activity, whereas EthD-2 fluorescence (red) identifies cells with compromised plasma membrane integrity. Increasing JH-RE-06 concentrations were associated with reduced Calcein-AM fluorescence, increased EthD-2 fluorescence, and changes in organoid morphology. Scale bars, 500 µm.

